# Incomplete Reprogramming of DNA Replication Timing in Induced Pluripotent Stem Cells

**DOI:** 10.1101/2023.06.12.544654

**Authors:** Matthew M. Edwards, Ning Wang, Dashiell J. Massey, Dieter Egli, Amnon Koren

## Abstract

Induced pluripotent stem cells (iPSC) are a widely used cell system and a foundation for cell therapy. Differences in gene expression, DNA methylation, and chromatin conformation, which have the potential to affect differentiation capacity, have been identified between iPSCs and embryonic stem cells (ESCs). Less is known about whether DNA replication timing – a process linked to both genome regulation and genome stability – is efficiently reprogrammed to the embryonic state. To answer this, we profiled and compared genome-wide replication timing between ESCs, iPSCs, and cells reprogrammed by somatic cell nuclear transfer (NT-ESCs). While NT-ESCs replicated their DNA in a manner indistinguishable from ESCs, a subset of iPSCs exhibit delayed replication at heterochromatic regions containing genes downregulated in iPSC with incompletely reprogrammed DNA methylation. DNA replication delays were not the result of gene expression and DNA methylation aberrations and persisted after differentiating cells to neuronal precursors. Thus, DNA replication timing can be resistant to reprogramming and lead to undesirable phenotypes in iPSCs, establishing it as an important genomic feature to consider when evaluating iPSC lines.

## Introduction

Reprogrammed pluripotent stem cells are widely used in research and provide the foundation for cell therapy (Bragança et al., 2019; Shi et al., 2017). Stem cells derived via somatic nuclear transfer to an enucleated oocyte (NT-ESCs) appear to be fully reprogrammed to the embryonic state and both mouse and human NT-ESCs can be reliably differentiated to specialized cell types (Kim et al., 2010; Ma et al., 2014; Sui et al., 2018). However, they are difficult to derive, in part because it requires access to human oocytes and highly specialized expertise. In contrast, induced pluripotent stem cells (iPSCs) can be readily derived by expressing defined factors in somatic cells, however their use is complicated by substantial variation in differentiation potential among iPSC lines and compared to their ESC counterparts (Koyanagi-Aoi et al., 2013; Sui et al., 2018). For example, some iPSC lines fail to undergo complete neural differentiation and give rise to teratomas following transplantation (Koyanagi-Aoi et al., 2013). Similarly, iPSCs show highly variable and reduced ability to differentiate to beta cells compared to ESC or reprogrammed NT-ESC (Sui et al., 2018). This raises important questions regarding the molecular basis of these differences, as well as regarding quality controls required for both research and cell therapy.

iPSC differentiation potential in both humans and mice is impacted by the incomplete reprogramming of primary cells to the stem cell state (Kim et al., 2010; Lister et al., 2011; Ma et al., 2014; Marchetto et al., 2009; Ohi et al., 2011). Studies in mice have reported that epigenetic memory can cause iPSCs to maintain epigenetic hallmarks of their cells-of-origin, predisposing them to differentiate back to the original cell type (Bar-Nur et al., 2011; Kim et al., 2010; Polo et al., 2010). Mouse iPSCs derived from beta cells, for example, show a propensity to differentiate to insulin producing cells (Bar-Nur et al., 2011), while fibroblast-derived iPSCs are resistant to differentiation into non-fibroblast lineages such as haematopoietic cells (Kim et al., 2010). This effect of source cell type on differentiation capacity, however, has not been observed in human cells.

Another possible source of variation in iPSCs are aberrant epigenetic changes acquired during the reprogramming process itself. These are typically sites of DNA methylation that are not seen in somatic donor cells or in ESCs (Lister et al., 2011; Panopoulos et al., 2017; Ruiz et al., 2012; Stadtfeld et al., 2010), and that can adversely affect developmental potential. These aberrations are both non-random and generally consistent across studies (Panopoulos et al., 2017). The most common form of such differentially methylated regions (DMRs) between iPSCs and ESCs are short (1-11Kb) regions that impact cytosines in the context of CpG dinucleotides (CG DMRs). At the majority of these DMRs, iPSCs are hypomethylated compared to ESCs. A second type of DMRs involve megabase-scale non-CG sites (methylation at CH sites, where H = A, T, or C) (non-CG DMRs)(Lister et al., 2011). Ultimately, both epigenetic memory and aberrant changes due to reprogramming have been reported at many genomic loci (Lister et al., 2011; Panopoulos et al., 2017), and deserve attention when considering epigenetic variation in iPSCs.

In addition to epigenetic alterations in iPSC lines, gene expression alterations have been characterized, with genes such as TCERG1L, FAM19A5, and COL22A1 reported across several studies to be differentially expressed between iPSCs and ESCs (Koyanagi-Aoi et al., 2013; Kyttälä et al., 2016; Lister et al., 2011). Such gene expression alterations, and the general heterogeneity of gene expression in iPSCs, pose challenges for using iPSCs for disease modeling (Rouhani et al., 2014). Similarly, some genes, such as TCERG1L, show consistent DNA methylation differences between ESCs and iPSCs (Koyanagi-Aoi et al., 2013; Lister et al., 2011). Another property that has been shown to resist reprogramming to the ESC state is chromosome conformation: some iPSCs derived from neuronal precursor cells (NPCs) fail to establish ESC-like chromosomal conformations on a small (sub-Mb) scale (Beagan et al., 2016), although topologically associating domains (TADs) were faithfully reconstructed in iPSCs (Krijger et al., 2016).

Notwithstanding these proposed molecular differences between iPSCs and ESCs, It remains unclear whether they fully account for the reduced differentiation potential of iPSCs. While expression alterations between iPSCs are frequently observed, it has been consistently shown that gene expression differences between iPSCs due to cell-of-origin (epigenetic memory) are subtle compared to differences brought about by inter-individual genetic variation (Burrows et al., 2016; Guenther et al., 2010; Kilpinen et al., 2017; Koyanagi-Aoi et al., 2013; Kyttälä et al., 2016). Aside from the handful of well-described genes, differential gene expression varies substantially between studies, suggesting that consistent reprogramming-associated gene expression alterations may be limited. Similarly, it has been shown that DNA methylation differences in iPSCs compared to ESCs, as well as between iPSCs from different primary cell types, are more subtle than those resulting from interindividual genetic variation (Burrows et al., 2016; Choi et al., 2015). Clustering based on either gene expression or DNA methylation groups cell lines by donor (genetic background) and fails to separate iPSCs from ESCs (Bock et al., 2011; Johannesson et al., 2014; Koyanagi-Aoi et al., 2013; Kyttälä et al., 2016). The insufficiency of gene expression and DNA methylation aberrations in accounting for the differentiation potential of iPSCs raises the question of whether other epigenetic properties may underly iPSC differences from ESCs.

A less explored epigenetic property that must undergo genome-wide changes as somatic nuclei dedifferentiate back to a stem-cell-like state is DNA replication timing. The coordinated replication of the genome throughout S-phase through the firing of replication origins is a highly regulated process unique to a given cell type (Hansen et al., 2010; Ryba et al., 2010). However, our knowledge of the ability of cells to reprogram DNA replication timing, and the differences in DNA replication timing between ESCs and reprogrammed cell types such as iPSCs and NT-ESCs, remain limited.

Examination of DNA replication dynamics with florescent *in situ* hybridization (FISH) has shown that both iPSCs and NT-ESCs fully reprogram DNA replication timing at several developmentally regulated loci (such as Rex1) as well as imprinted genes (such as SNRPN and IGF2R) (Shufaro et al., 2010). However, this approach is limited both in its genomic resolution (specific loci) and its temporal resolution. Genome-scale analysis of iPSC reprogrammed from fibroblasts has previously suggested that their DNA replication timing is “virtually indistinguishable” from ESCs (Hiratani et al., 2008; Ryba et al., 2010), however this analysis was done at the level of large replication “domains” and on a small number of predominantly mouse cell lines. In contrast, a more recent study using a single molecule approach has reported incomplete reprogramming of DNA replication timing at the FXN and NANOG loci in iPSCs, but not NT-ESCs (Paniza et al., 2020), although only three loci were examined with a small sample size. A high-resolution, genome-wide analysis of iPSC and NT-ESC DNA replication timing compared to ESC is currently lacking, and necessary to resolve replication dynamics in reprogrammed stem cells.

Here, we generated and compared genome-wide DNA replication timing profiles of multiple ESC, iPSC, and NT-ESC lines. While NT-ESCs faithfully recapitulated the DNA replication timing program of ESCs, we identified 26 genomic regions with altered replication timing in a subset of iPSCs, most of them replication delays. iPSC replication timing aberrations were located primarily in centromere- and telomere-proximal regions and were enriched for H3K9me3 and zinc finger repeats. These regions were related to previously identified DMRs and genes with altered expression in iPSCs, though the replication timing alterations appear distinct from these other two epigenetic properties. Finally, we demonstrate that replication timing alterations persist in NPCs derived from aberrantly replicating iPSCs, which carries important implications for their use in research and cell therapy.

## Results

### Aberrant DNA replication timing in iPSCs

To comprehensively compare the DNA replication timing program across pluripotent stem cells derived by different methods (**Fig. 1A**), we concomitantly profiled genome-wide DNA replication timing in 28 ESC lines, four NT-ESC lines, and 19 iPSC lines (Supplementary Table 1). Of the four NT-ESCs, three (NT5, NT6 and NT8) were derived from a neonatal fibroblast cell line (BJ) and one was derived from adult dermal fibroblasts (1018); two iPSCs were also derived from neonatal BJ cells and two from adult 1018 cells, allowing for isogenic NT-SC/iPSC comparison. We also profiled neuronal precursor cell lines derived from either ESCs or iPSCs to examine differentiation in different stem cell types (see further below).

**Figure 1:**
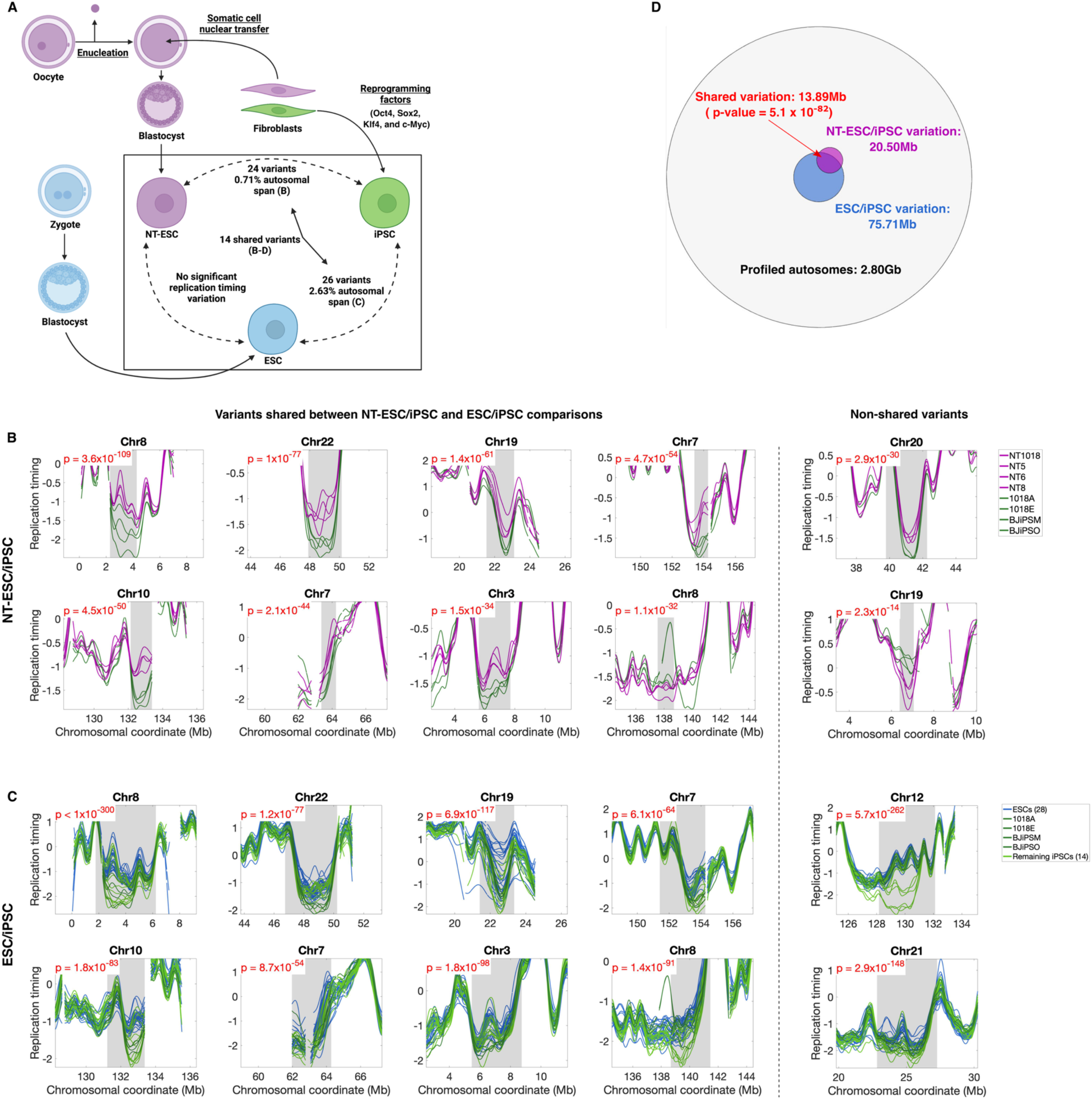
NT-ESCs, but not iPSCs, faithfully reprogram replication timing. **A.** Derivation of NT-ESCs, iPSCs, and ESCs. Box: summary of DNA replication timing differences between stem cell types. iPS cell lines were created using the standard four Yamanaka factors. Panel created with Biorender.com. **B.** Four NT-ESCs (purple) were compared to isogenic iPSC (green) in sliding windows along the genome using ANOVA. Ten of 26 variants are shown (the remainder are shown in **Fig. S2A** and **B**), including the eight most significant variants of the 14 which overlapped ESC/iPSC variants (left), and the two most significant of the remaining 10 unshared variants (right). **C**. iPSCs (green) were compared to ESCs (blue) using an empirical ANOVA scan at a 5% FDR cutoff (**Methods**). The eight variants on the left are the same as those shown in **B**. Windows shown are centered around the ESC/iPSC variants with +/-3Mb flanking on either side. As these were the primary variants we considered (see below), the variants in **B** were aligned to these same coordinates. The most significant of the remaining 12 variants are shown on the right. The remaining variants are shown in **Fig. S2A** and **C**. **D.** A Venn-diagram of ESC/iPSC and NT-ESC/iPSC variation compared with the rest of the profiled autosomes. Overlap between ESC/iPSC variants and NT-ESC/iPSC variants was significant when compared to 1000 permutations (p = 5.1 x 10^-82^; **Methods**).

Replication timing was profiled at high-resolution using a whole genome sequencing (WGS)-based approach which we have previously described (Koren et al., 2014, 2021). Briefly, earlier-replicating regions of the genome contribute more DNA in a population of proliferating cells than do later-replicating regions, which can be observed in the form of fluctuating DNA copy number along chromosomes in WGS data. Accordingly, we calculated the number of sequencing reads in uniquely-alignable genomic windows, corrected for the influence of GC content, and filtered copy number variations and outliers. The remaining DNA copy number profiles were smoothed and normalized to standard deviation units to derive the replication timing profiles of each cell line and compare them across samples. The replication timing profiles were of high quality, as evidenced by high correlations within ESCs (r = 0.96), iPSCs (r = 0.97), and NT-ESCs (r = 0.98). The replication profiles of all cell lines also resembled previously profiled ESCs (Ding et al., 2021) (mean r = 0.84), more so than they resembled a different cell type (lymphoblastoid cell lines, mean r = 0.71). One iPSC line was removed based on correlation to other iPSCs and principal component analysis (**Fig. S1A,B**).

Correlations between stem cell types (ESC/NT-ESC: r = 0.97; ESC/iPSC: r = 0.96; NT-ESC/iPSC: r = 0.97) were comparable to those within stem cell types, indicating that globally, replication timing is consistent between reprogrammed and embryonic stem cells. Nonetheless, we observed notable differences in the replication profiles at the local scale. To quantify this systematically, we first compared NT-ESCs and their isogenic iPSC counterparts to identify genomic regions where replication timing differed between reprogramming methods. We used an approach we employed previously (Edwards et al., 2021), where candidate regions were first identified using ANOVA tests in approximately 200Kb-wide overlapping windows, and then were filtered to remove regions unlikely to contain true variation (see **Methods**). We thus identified 24 regions of replication timing variation (**Fig. 1B; Fig. S2A, B**).

To further understand the source of this variation, we compared cell lines derived by both reprogramming methods to ESCs. Because of the larger number of samples available for this analysis, we used permutations to empirically set a p-value threshold corresponding to a 5% false discovery rate (FDR; **Methods**). This identified 26 ESC/iPSC variants, covering a total of 2.63% of the autosomes (**Fig. 1C; Fig. S2A, C**), at a p-value threshold of 1 x 10^-53^. We found that these ESC/iPSC variants had a significant and considerable degree of overlap with NT-ESC/iPSC variation (14/26, or 53.8% of variants overlapped, p = 5.2 x 10^-82^; **Fig. 1D**). We also directly tested NT-ESC/iPSC variants using all 18 iPSCs (not just isogenic lines) and the genome-wide Bonferroni-corrected cutoff from the filtering-based ANOVA scan, and confirmed that 15/24 (62.5%) variants were identified in this larger set (**Fig. S3A**). In contrast, the ESCs/NT-ESC comparison did not reveal any variation at a 5% FDR. A down-sampling analysis confirmed that the smaller sample size when examining NT-ESCs vs isogenic iPSCs did not bias these results (**Methods**). Consistent with iPSCs being the outlier stem cell type, the 18 iPSCs showed lower correlation to NT-ESCs at variant regions (r = 0.93) compared to non-variant regions (r= 0.99), while ESCs were highly correlated (r = 0.99) to NT-ESCs at both variant and non-variant regions (**Fig. S3B**). Given the high correspondence between NT-ESCs and ESCs, and the larger number of ESC samples, we focused our subsequent analyses on replication timing differences between ESCs and iPSCs.

Notably, 22/26 (84.6%) of the ESC/iPSC variants consisted of replication delays in iPSCs, with mean iPSC replication timing later than the means of both ESCs and NT-ESCs (**Fig. 1B-C, Fig. S2D**). Replication was already late in ESCs in these regions, with an average replication timing of 0.73 SD units later than the mean, while iPSCs replicated even later, at an average of 0.91 SD units later than the mean (**Fig. S2E**). Many of these regions showed prominent replication timing peaks in ESCs, indictive of replication origins or origin clusters. These peaks appeared diminished or even lost in some iPSCs (see for examples variants on chromosome 10, 12, and the strongest variant on chromosome 8 in **Fig. 1C**), suggesting that replication origin firing may be inhibited in iPSCs.

To further validate the ESC/iPSC replication timing variants that we identified, we turned to a larger, independent dataset of replication timing in 108 ESCs and 300 iPSCs (Ding et al., 2021). We were able to test 17 of the 26 variants in this dataset (**Methods**) and found that the iPSCs in the larger dataset showed the same direction of variation (delayed/advanced) in accordance with expectation at 14 of 17 variants (82%). Furthermore, nine of the 17 testable variants (53%) showed similar replication timing alterations as those initially observed in the smaller dataset (**Fig. S4; Methods**). We did not attempt to identify new variants using these larger sample sizes because of concerns of batch effects, as the two cell types were sequenced in separate projects.

Taken together, these results suggest that NT-ESCs closely resemble ESCs in their replication dynamics, while iPSCs show replication timing alterations, largely consisting of delays, at specific genomic regions when compared to either type of ESCs.

### Replication timing delays at TCERG1L

One of the most significant variant region we identified in both the ESC/iPSC and NT/iPSC comparisons (Chr10:131,222,871-133,380,323; **Fig. 1B-C**) contained the gene TCERG1L, identified in several previous studies as having differential methylation and decreased gene expression in iPSCs compared to ESCs (Lister et al., 2011; Nishino et al., 2011; Ruiz et al., 2012). Specifically, iPSCs replicated later than ESCs at this locus, consistent with decreased gene expression. This is in contradiction with the results of Single Molecule Analysis of Replicated DNA (SMARD), which suggested similar replication dynamics between ESCs and iPSCs at TCERG1L (Paniza et al., 2020). SMARD also suggested incomplete replication timing reprogramming of the FXN and NANOG loci, however neither locus showed significant variation in our data between the two pluripotent cell types (p = 0.77 and 0.045 respectively, compared to p = 1 x 10^-53^ required to achieve genome-wide significance as detailed above; **Fig. S5A**).

Similar results were observed when testing the exact cell lines used in the previous study (the four NT-ESCs and their isogenic iPSCs; **Fig. S5B**). SMARD also reported that iPSCs derived from adult cells (1018A/E) have lower initiation frequency at the FXN locus than those derived from neonatal cells (BJiPSM/BJiPSO), however we did not observe this either (**Fig. S5B**). We ascribe these inconsistencies to the different methodologies and the limited genomic scale of analysis, as well as the limited number of samples and fibers used by SMARD.

### Replication timing alterations identify a subset of aberrant iPSC lines

While replication was altered in iPSCs, there appeared to be substantial heterogeneity in replication timing alterations among individual iPSC lines. Specifically, in several variant regions some iPSC lines resembled ESC replication timing while others had delayed replication. To quantify this heterogeneity, we compared replication timing for each ESCs and iPSCs cell line at both variant and non-variant regions. We observed high correspondence in replication timing at non-variant regions for both iPSCs (correlation to the ESC mean ranged from r = 0.96 to 0.99) and ESCs (correlation to the iPSC mean ranged from r = 0.95 to 0.99). At variant regions, iPSCs and ESCs line showed slightly lower correspondence, but were nonetheless still similar to each other (median r = 0.91 and 0.90 to the ESC and iPSC means, respectively). However, iPSCs exhibited much greater variability (correlation to the ESC mean ranged from r = 0.66 to 0.97) compared to ESCs (correlation to the iPSC mean ranged from r = 0.81 to 0.94); **Fig. 2A**), suggesting heterogeneity among iPSCs at variant regions.

**Figure 2.**
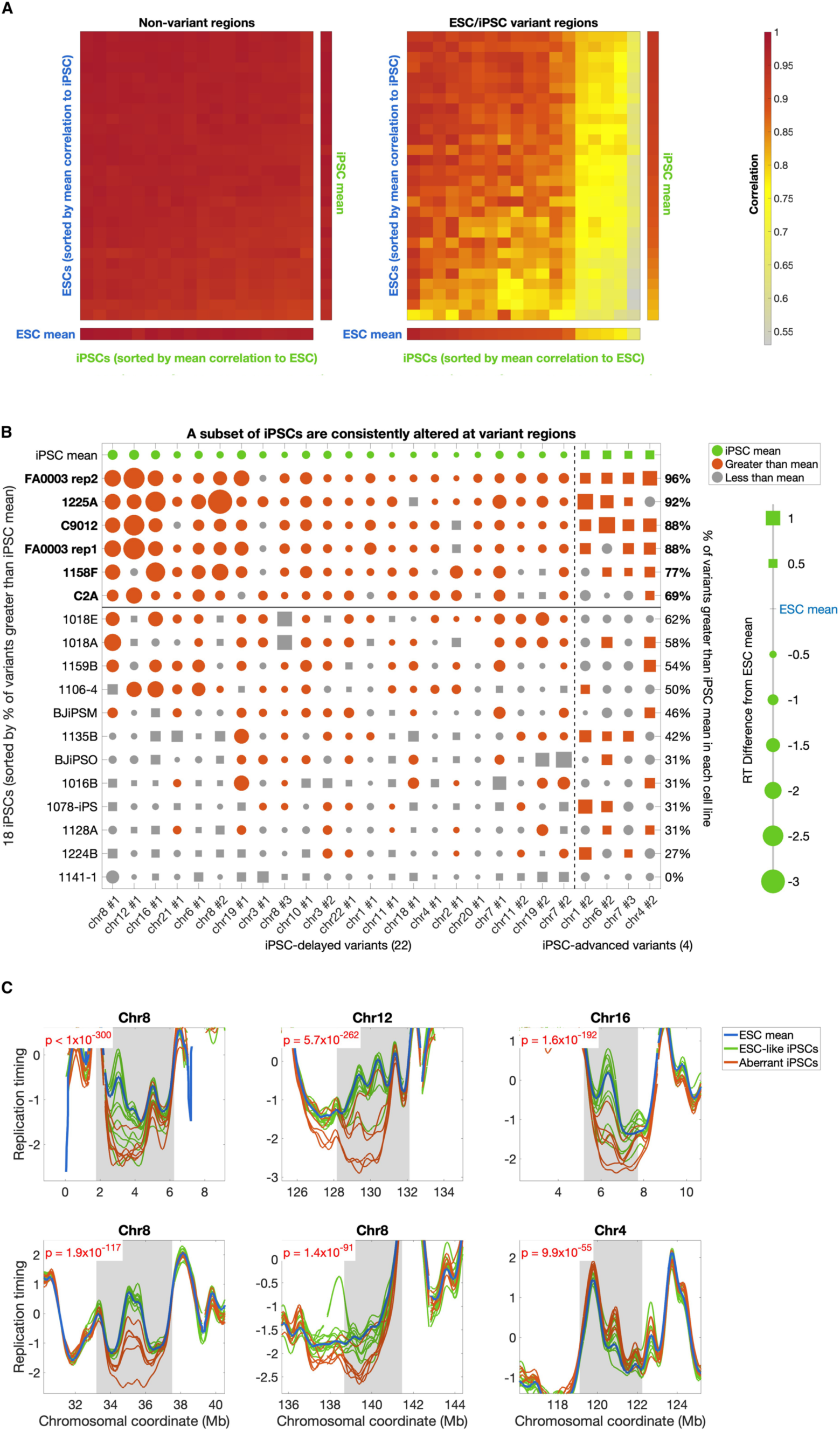
Heterogenous replication delays in iPSCs. **A**. Correlation of replication timing between ESCs (rows) and iPSCs (columns) at non-variant regions (left) and at ESC/iPSC variant regions (right). **B.** Replication timing difference from the ESC mean for each individual iPSC line. Values are shown as squares for replication timing earlier than the ESC mean, and circles for replication timing later than the mean; the size of each marker reflects the extent of advance/delay at the region of maximum variation (scale to the right). Replication timing is shown for each of the 26 variant regions, first separated into iPSC delayed (left) and iPSC advanced (right), and then ordered from left to right by significance (based on the p-values from Fig. 1C and **S2A,C**; variants are labeled by chromosome first and then numbered in increasing p-value order; i.e., chr8 #2 is the second lowest p-value variant on chr8). The iPSC mean (green) is shown in the top row, and values for each cell line are indicated as more variant than the mean (orange; e.g., later than the mean at a delayed variant), or less variant (grey; e.g., earlier than the mean at a delayed variant). Cell lines are ordered by the percentage of regions which are more variant than the mean (values on the right). The six cell lines which were more variant than expected by chance (binomial test, see text) are indicated in bold and classified as aberrant iPSCs, while the remaining twelve were classified as ESC-like. **C.** Six example variant regions, with the six aberrant iPSCs shown in orange. The first five variants were iPSC-delayed, while the bottom right variant is iPSC-advanced.

To determine if this heterogeneity originated from across iPSC lines or resulted from a subset of them, we quantified the extent of replication timing aberration in each individual iPSC line, at the region of maximum variation (maximum absolute replication timing difference between each iPSC and the ESC mean) in each variant (**Fig. 2B**). This separated the iPSC lines into two groups: those that resembled the replication timing of ESCs, and others that were altered at nearly every variant. Accordingly, we identified the iPSCs that were more aberrant than the iPSC mean at each variant (more delayed at iPSC-delayed variants, more advanced at iPSC-advanced variants), and defined those that were consistently more aberrant in at least 18 of 26 variants (>= 69%; binomial p = 0.038) as “aberrant iPSCs”. The six cell lines thus identified had similar replication timing alterations at each variant region (**Fig. 2C)**. A genome-wide scan for replication timing variation (empirical ANOVA scan, 5% FDR) between the remaining 12 iPSCs and ESCs did not identify any significant variation, indicating that variation was driven primarily by the aberrant iPSCs. Accordingly, we term the remaining iPSCs as “ESC-like”.

We also observed heterogeneity among individual iPSC (but not ESC) lines in the larger dataset of 300 iPSC and 108 ESCs that we used above for validation (**Fig. S6A**). Once again, we were able to identify two opposing subsets of iPSC lines based on the correlation of each cell line’s replication profile to ESCs and the maximum replication timing alteration (across the seven iPSC-delayed regions validated in this dataset; **Fig. S4**) relative to ESCs. Using a cutoff of the 10% most outlying cell lines in both of the above criteria, we defined 16 aberrant iPSCs and 14 ESC-like iPSCs (**Fig. S6B**). When comparing replication timing between these two groups, there were clear differences at variant regions, with the ESC-like iPSCs largely resembling ESCs and the aberrant cell lines showing replication delays (**Fig. S6C**).

### iPSC replication timing variants are associated with repressed genes and heterochromatin

Interestingly, 31% (8/26) of ESC/iPSC variants were located in the 10% of chromosome arms closest to telomeres (telomere-proximal), while another 19% (5/26) were located in the 10% closest to centromeres (centromere-proximal) (**Fig. 3A,B**). Since both centromere- and telomere-proximal regions are heterochromatic (Benetti et al., 2007; Gonzalo et al., 2006; Saksouk et al., 2015), we separately characterized epigenetic features of those variants proximal to centromeres or telomeres, as well as the remaining variants. We tested for enrichment of 31 histone marks as well as DNase hypersensitivity (Hoffman et al., 2013) and chromatin states defined by ChromHMM (Ernst & Kellis, 2017). The histone mark H3K9me3, an established marker of constitutive heterochromatin, was greatly enriched specifically in telomere-proximal variants (**Fig. 3C**), while H3K27me3, a marker of facultative heterochromatin, was not. Centromere-proximal variants were also significantly depleted of the active mark H3K4me1 (**Fig. 3C**). The remaining variant regions that were not near centromeres or telomeres did not show any histone mark enrichment (**Fig. S7A**). The ChromHMM state “heterochromatin” was found to be enriched in all three classes of variants (**Fig. 3D, Fig. S7B**), while there was a substantial enrichment of zinc finger (ZNF) genes & repeats in centromere-proximal regions (**Fig. 3D**). Although the statistical enrichment of ZNF genes & repeats was largely due to two ZNF clusters at the two variants on either side of the chr19 centromere (**Fig. 1C and S2A**), this chromatin state was found in all five centromere-proximal variants.

**Figure 3:**
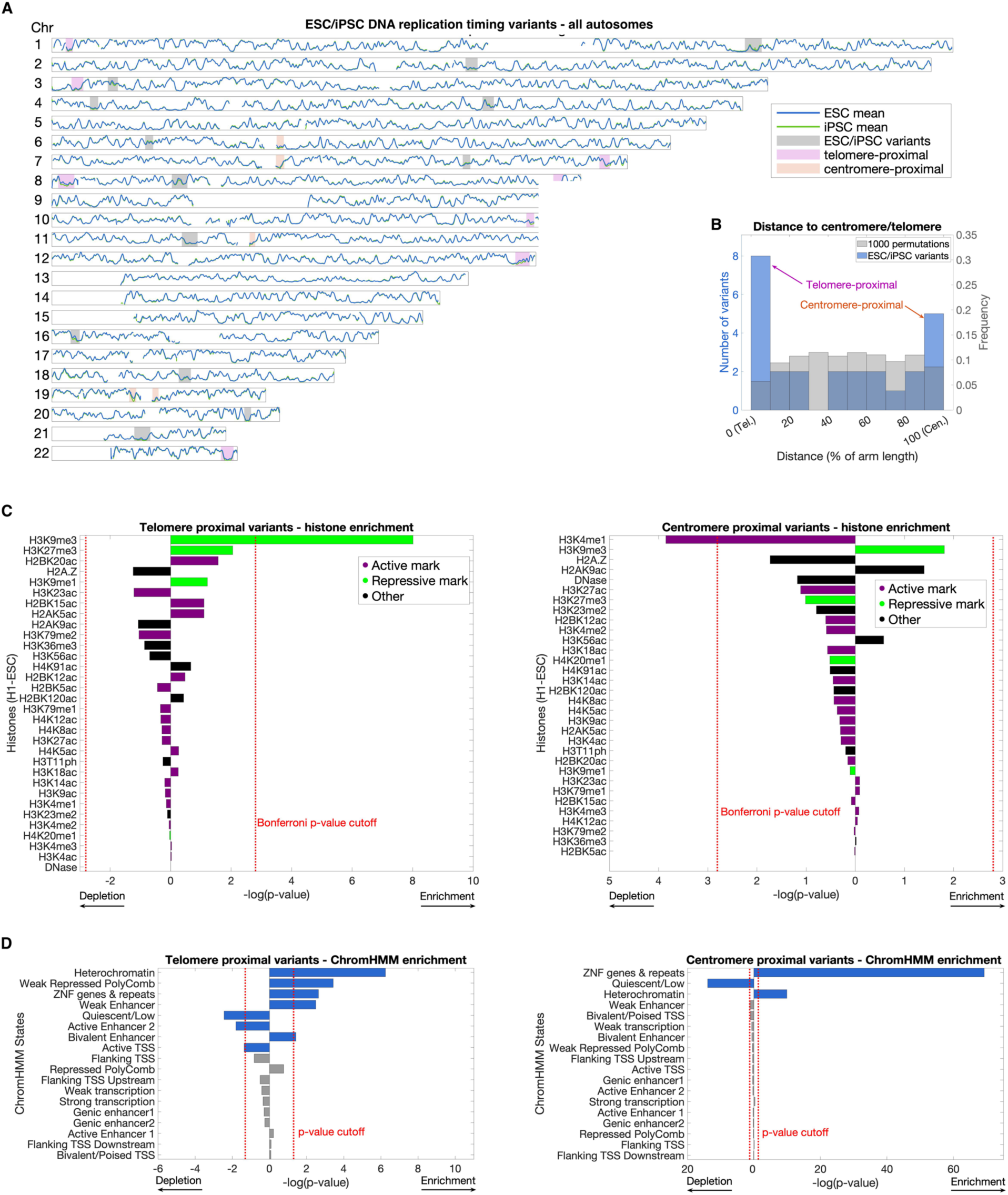
Replication timing variants are located in heterochromatic regions near centromeres and telomeres. **A.** Whole genome plot of the mean ESC and iPSC replication timing profiles. Variant regions are shaded, and variants within 10% of chromosome arm length of telomeres (purple) and centromeres (orange) are indicated. **B.** Histogram of the relative locations along the chromosome arm of the 26 variants (blue; left y scale) compared with 1000 random permutations of variant region locations across the genome (grey; right y scale). Distance is calculated between the middle of each variant and the nearest telomere and normalized by the length of the chromosome arm. Variants within 10% of telomeres or centromeres were considered as centromere- or telomere-proximal, respectively. **C.** Analysis of histones in telomere-proximal variants (left) and centromere-proximal variants (right) based on ChIP-seq data (ENCODE). Bars indicate enrichment/depletion two-sided t-test p-value of each chromatin mark (purple: active marks, green: repressive marks) in variants compared to 1000 replication timing matched permutations. Each chromatin mark was considered to be independent, and the Bonferroni corrected p-value threshold (.05/32) is shown in red. Telomere-proximal variants are significantly enriched for the histone mark H3K9me3, while centromere-proximal variants are nominally enriched for H3K9me3 and significantly depleted of H3K4me1. **D.** Same as **C** for ChromHMM states in telomere-proximal variants (left) and centromere proximal variants (right). Because these states are by definition mutually exclusive, we considered a cutoff of p = 0.05 for significance (red line). ChromHMM state counts were defined by the number of base pairs belonging to each state in variant regions and permutations. Both sets of variants were enriched for heterochromatin and depleted for quiescent regions.

There were 202 genes located within the variant regions, in line with what is expected for similarly late-replicating regions (p=0.64 when compared to 1000 permutations; **Methods**). Gene ontology analysis revealed a single enriched category, “*cis*-regulatory region sequence specific binding”, which was predominantly attributed to a cluster of zinc finger protein-encoding genes at a single variant (3.23-fold enrichment, FDR = 0.0282, 15 of 18 genes located at chr19:21,405,159-23,345,945). Among these genes were FAM19A5 (TAFA5), FZD10, TMEM132D, COL22A1, and TCERG1L (**Fig. 4C**), which have all been shown to be differentially expressed in iPSCs (Kyttälä et al., 2016; Liang & Zhang, 2013; Lister et al., 2011; Ruiz et al., 2012).

**Figure 4:**
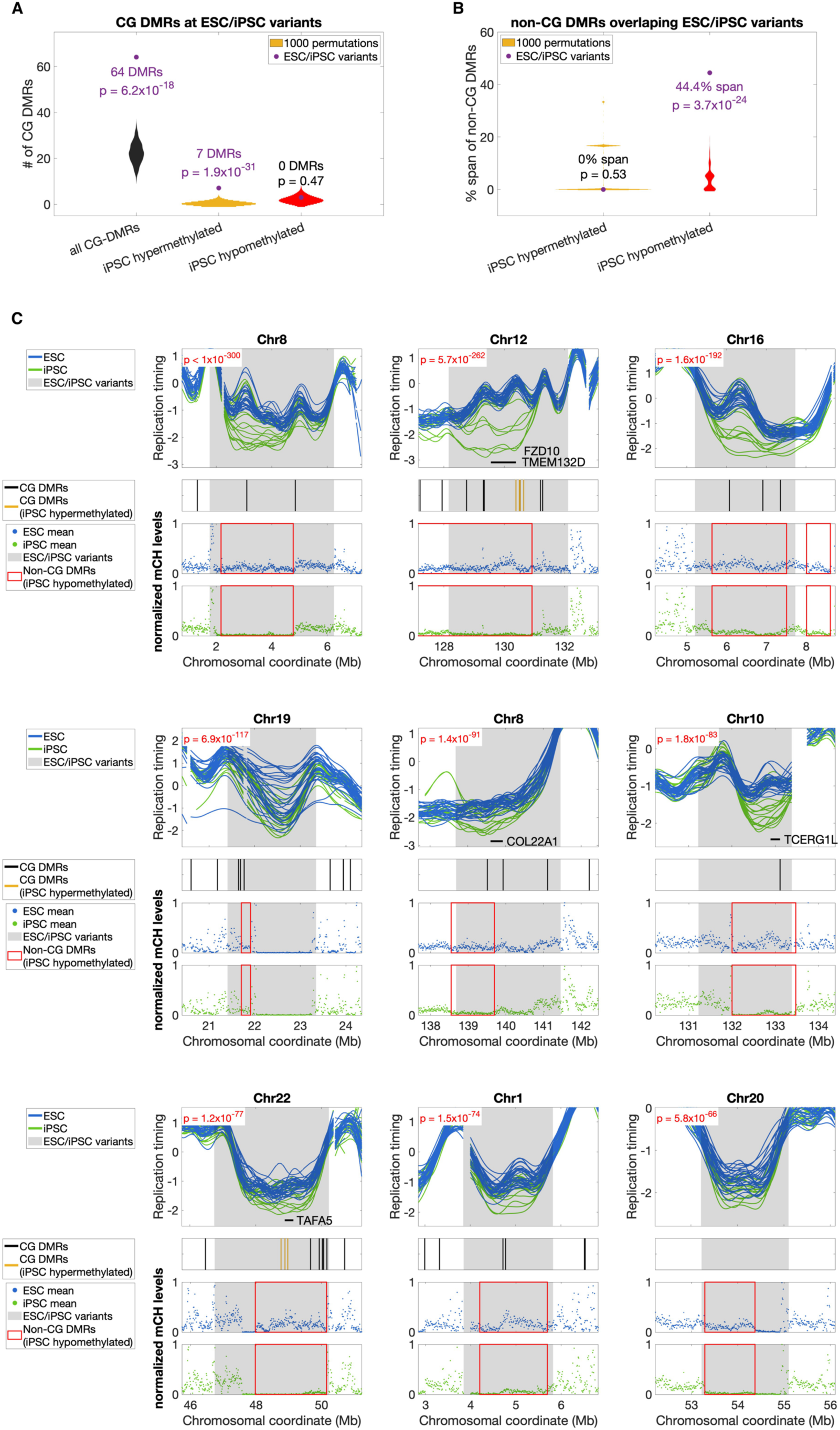
Regions of iPSC replication timing variation correspond to differential CG and non-CG DNA methylation. **A.** Distributions of CG DMRs in 1000 permutations of ESC/iPSC variant regions (Violin plots; see **Methods**). Distributions were smoothed using the ksdensity MATLAB function with robust kernel estimation (via the DistributionPlot function). The total number of the CG DMRs in variant regions is represented as a purple dot; two-tailed p-value calculated from z-score compared to the distribution is indicated (purple font indicates a significant p-value). **B.** Similar to **A**, for regions of non-CG hypomethylation in iPSCs. Because all non-CG DMRs were either hypermethylated or hypomethylated (unlike in **A** where most DMRs were unclassified), only these two sets of DMRs were compared to variant regions. Additionally, due to the larger size of these non-CG DMRs, overlap was calculated as cumulative percentage of each replication timing variant spanned by the DMR (see **Methods**). Because the iPSC hypermethylated non-CG DMRs (orange) were on average smaller than replication timing variants (mean sizes of 526Kb and 2.92Mb, respectively), the distribution of overlaps follows several discrete modes. The hypomethylated non-CG DMRs (red; mean size of 1.38Mb) were larger, and thus the distribution of overlaps was smoother. **C.** The nine ESC/iPSC variant regions that overlapped a non-CG iPSC hypomethylated regions are shown with corresponding differentially expressed genes (DEGs) and DNA methylation data. iPSC CG DMRs are shown as vertical black lines (and are enriched in variant regions, see **A**), while those consistently hypermethylated in iPSCs are indicated in orange. Non-CG methylation levels (ranging from 0 to 1 after normalization) are plotted for the mean of two ESCs (blue) and the mean of five iPSCs (green); red boxes: non-CG DMRs hypomethylated in iPSCs (see **Methods** for methylation data processing)

### Replication timing variation in iPSCs is related to DNA methylation aberrations

It has previously been shown that DNA methylation is altered in iPSCs compared to ESCs due to both incomplete reprogramming to the stem cell state at certain genomic regions, and aberrant methylation at other regions (Lister et al., 2011). We thus asked whether replication timing variants are associated with regions of differential methylation. Lister et al. identified 1,175 short CG-DMRs between ESCs and iPSCs, among which 11 were consistently hypermethylated in all iPSCs and 119 were consistently hypomethylated (the remainder were not consistent across tested cell lines). In addition to these short CG-DMR, Lister et al. identified 29 long non-CG DMRs, seven of which were hypermethylated in iPSCs relative to ESCs, while the remaining 22 were hypomethylated and considered not fully epigenetically reprogrammed. Interestingly, non-CG DMRs were enriched near centromeres and telomeres, similar to replication timing variants.

Sixty-four (5.45%) of the CG-DMRs overlapped our ESC/iPSC replication timing variant regions, significantly more than expected by chance (p = 6.2 x 10^-18^; **Fig. 4A,C**). Notably among those, seven of the 11 (63.6%) hypermethylated CG-DMRs overlapped replication timing variants, a strong enrichment compared to chance (p = 1.9 x 10^-31^; **Fig. 4A,C**), while there was no enrichment for the more abundant hypomethylated CG-DMRs in variant regions (p = 0.47; **Fig. 4A**). In contrast to CG-DMRs, hypomethylated non-CG DMRs showed a strong correspondence to the ESC/iPSC variants (p = 3.7 x 10^-24^; **Fig. 4B,C**), while hypermethylated non-CG DMRs did not overlap with replication timing variation (p = 0.54; **Fig. 4B**). Specifically, of 19 non-CG DMR hypomethylated regions that we were able to test (successfully lifted over to hg19), nine (47%) at least partially overlapped nine of our 26 variant regions (**Fig. 4B-C**), all of which were iPSC-delayed. Moreover, directly testing these regions revealed that 14/19 (73.7%) DMRs (including eight that overlapped ESC/iPSC variant regions) showed ESC/iPSC replication timing variation that was significant at a genome-wide Bonferroni-corrected p-value threshold of 8.63 x 10^-7^ (**Fig. S8**; note that six of these were not identified as replication timing variants in our original scan because the individual constituent windows were not significantly different). Taken together, replication timing differences between iPSCs and ESCs correspond to aberrant methylation in iPSCs involving both short, hypermethylated CG DMRs and long, hypomethylated non-CG DMRs.

### Aberrant replication timing in iPSCs persists after neuronal differentiation

The replication timing alterations we identified suggest that some iPSCs may be incompletely reprogrammed or have newly acquired epigenetic aberrations; this observation is important for evaluating the utility of such iPSC lines, for instance as models for cellular differentiation. To test whether iPSC replication timing alterations persist after differentiation, we differentiated three iPSC lines that were identified as aberrant and three that were ESC-like, as well as eleven ESC lines, to NPCs (**Fig. S1C**), and profiled replication timing together with the source stem cell lines. Two cell lines derived from ESC, and one from an ESC-like iPSC were removed based on correlation to other NPCs and principal component analysis (**Fig. S1A,B**).

As expected for different cell types, undifferentiated and differentiated cells were less correlated with each other (r = 0.82) compared to the correlations among cell lines within each differentiation state (ESC: r = 0.96, iPSC: r = 0.97, NPC: r = 0.96) or compared to the correlations between ESCs and iPSCs (r = 0.96; **Fig. S1A**).

Correlations among ESC-derived NPCs (r = 0.97), among iPSC-derived NPCs (r = 0.95) and between the two groups of NPCs (r = 0.96) were similar, suggesting that the type of stem cells used to derive NPCs has little impact on the global replication timing program. ESC-derived and iPSC-derived NPCs also appeared similar visually at the chromosomal scale (**Fig. 5A**). In contrast, at ESC/iPSC replication timing variant regions, we also observed variation in those NPCs derived from aberrant iPSCs (herein referred to as aberrant-derived NPCs; **Fig. 5A,B**). Systematic testing of replication timing variant regions using ANOVA revealed that aberrant-derived NPCs indeed showed variation compared to ESC-derived NPCs, which was both significant at a genome-wide Bonferroni-corrected significance threshold and in the same direction as the ESC/iPSC variation. Aberrant iPSCs were variant at 16/26 (58%) regions (**Fig. 5B**), an enrichment compared to permutations (9.7 x 10^-7^). We also called variation between ESC-derived and iPSC-derived NPCs using an unbiased genome-wide ANOVA scan, and found 27 regions of variation, spanning 3.2% of the autosomes, at a 5% FDR. Of these, nine (33.3%) overlapped nine ESC/iPSC variants (**Fig. 5B**; p = 2.6 x 10^-12^ when compared to 1000 permutations; **Methods**), suggesting that at least some of the replication timing alterations in iPSCs are maintained through differentiation. In certain cases, e.g., on chromosomes 12 and 16 (**Fig. 5**), replication delays appeared to become even more extreme after NPC differentiation.

**Figure 5:**
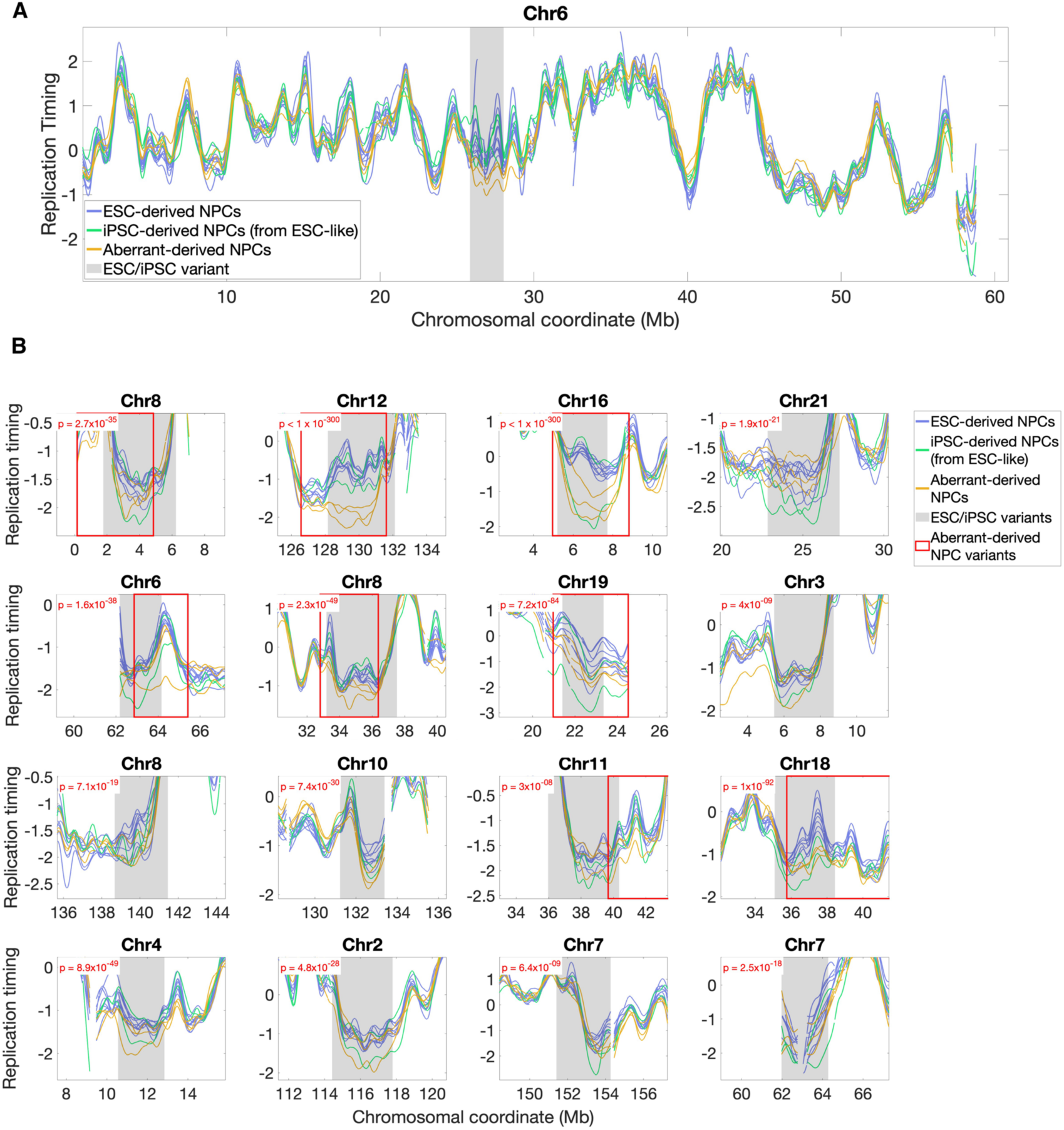
Replication timing aberrations persist after differentiation to NPCs. **A.** Replication timing profiles for ESC-derived NPCs (blue), ESC-like iPSC-derived NPCs (green), and aberrant-derived NPCs (orange) on the p arm of chromosome 6. An iPSC/ESC replication timing variant region (grey shaded area) is maintained through differentiation. **B.** Replication timing profiles for NPCs at the sixteen regions where aberrant-derived iPSCs were significantly different from ESC-derived NPCs at the genome-wide Bonferroni threshold (p = 9.58 × 10^−7^). Significant regions were also required to show the same direction of effect (delayed in all validated cases) as the ESC/iPSC variant (this removed two regions of variation).

## Discussion

Reprogrammed stem cells provide a necessary alternative to ESCs due to limitations on the ability to generate new ESC lines and their ability to be genetically and immunologically matched to individual patients. However, molecular variations involving genetic and/or epigenetic aberrations can greatly impact the use of iPSCs and exacerbate phenotypic variation including cells’ ability to differentiate to somatic cells of interest. Therefore, epigenetic deviations of reprogrammed cells compared to ESCs as well as genetic change compared to the somatic cell donor, raise concerns for both research and therapeutics. Here, we report that NT-ESCs provide a faithful recapitulation of the ESC DNA replication timing program. In contrast, iPSCs are prone to replication defects. We identify a total of 26 genomic regions, covering 2.63% of the autosomes, where a subset of iPSCs consistently deviate in their replication timing compared with ESCs. We validated these regions by comparing isogenic iPSCs and NT-ESCs, by comparing a larger set of iPSCs to NT-ESC, and by examining a yet larger dataset encompassing hundreds of ESC and iPSC cell lines. Evaluating replication dynamics across many cell lines controlled for natural and technical variation between samples, while a genome-wide analysis that considers the chromosomal vicinity of each locus controlled for any potential localized noise at specific sites; in these respects, our analysis is superior to previous methodologies and studies.

Given the tendency of ESC/iPSC variants to be near telomeres and centromeres, it is likely that we are missing additional variants in unmapped genomic regions including centromeres and telomeres proper. We also identified six additional variants when testing non-CG DMRs, suggesting that there may be additional variant regions we did not identify using our stringent genome-wide criteria. Similarly, there were 18 regions of variation specifically observed between aberrant-derived NPCs and ESC-derived NPCs, which could represent undiscovered ESC/iPSC variants which were only significant after differentiation, and/or regions at which replication timing variation emerged after differentiation.

Delayed DNA replication timing in iPSCs and their differentiated derivatives has important implications for their use, as it could lead to increase in mutation rates in the affected regions, to aberrant cell cycle progression, or possibly to alterations in genome regulation (Koren, 2014). Indeed, iPSCs are known to be prone to acquisition of somatic mutations, copy number alterations, and chromosomal rearrangements (Ben-David et al., 2011; Hussein et al., 2011; Lu et al., 2014; Pasi et al., 2011; Robinton & Daley, 2012), possibly due in part to delayed replication timing.

### Replication timing defects in iPSCs associate with, but are not fully explained by gene expression and DNA methylation alterations

iPSCs are known to exhibit altered gene expression, DNA methylation, and chromatin conformation compared to ESCs. Here, we have shown that iPSCs are also prone to alterations in the DNA replication timing program, which co-occur with, but are distinct from, these other epigenomic alterations. While several downregulated genes are located in regions of delayed replication in iPSCs, replication timing alterations are concentrated in late-replicating, gene-poor regions of the genome. These regions tend to be heterochromatic, enriched for H3K9me3, and are often localized near centromeres and telomeres. The larger number (at least 26) of replication timing variants compared to gene expression changes (only a handful consistent among studies) between ESCs and iPSCs suggests that DNA replication timing alterations are more abundant, span a greater fraction of the genome, and are at least partially independent of gene expression alterations. This raises important questions regarding the mechanism causing these differences, in particular the possibility of aberrant *de novo* changes that are neither present in the donor cells nor in embryonic stem cells.

Compared to gene expression changes, differences in DNA methylation in iPSCs relative to ESCs were more abundant at replication timing variant loci. Replication timing alterations coincided with both short CG-DMRs (particularly iPSC hypermethylation) and longer regions of non-CG hypomethylation in iPSCs. Previous studies have suggested that DNA replication timing is influenced by DNA methylation in some contexts but independent of it in other contexts (Du et al., 2019, 2021; Gribnau et al., 2003; Takebayashi et al., 2021). Consistently, we identified cases of replication timing alterations in iPSCs that occurred independently of DNA methylation aberrations, including near centromeres and telomeres.

### Cellular sources of altered DNA replication timing

Epigenetic alterations in iPSC compared to ESCs have been shown to arise from either epigenetic memory of the cells from which the iPSCs were derived (Bar-Nur et al., 2011; Kim et al., 2010; Polo et al., 2010), or alterations due to the reprogramming process (Lister et al., 2011). Similarly, abnormal replication timing in iPSCs could be due to a failure to sufficiently reprogram somatic cell (fibroblast in this case) replication timing, or aberrations due to reprogramming itself. Understanding whether replication defects are a remnant from fibroblasts, or a product of reprogramming could potentially be resolved by studying DNA replication timing in fibroblasts. However, fibroblasts proliferate significantly slower than stem cells and are not amenable to the WGS-based replication profiling utilized here. An alternative approach is to use single cell DNA sequencing data, for instance those available for the fibroblast cell line BJ (isogenic to NT5/6/8, BJiPSM, and BJiPSO; 10xgenomics.com), together with our recently published approach for inferring DNA replication timing from proliferating single cells (**Methods**; Massey & Koren, 2022). Such an analysis suggested that at four replication timing variants, iPSC replication resembled fibroblasts, possibly reflecting epigenetic memory of the source cell type; while at two other variants, fibroblasts resembled ESCs but not iPSCs, more consistent with a possible reprogramming defect in iPSCs (**Fig. S9**). The remaining variants were not informative, as fibroblasts did not resemble either stem cell type (see **Methods**; although this may itself be interpreted as refuting epigenetic memory). It is important to note that we cannot rule out coincidental similarity in either case given the small number of examples and the methodological differences. Thus, further research will be required in order to resolve the cellular sources of altered DNA replication timing in iPSC lines.

### Replication timing as a screen for differentiation potential

We have shown that reprogramming defects occur in a heterogeneous manner across cell lines, with variation being confined to a subset of iPSC lines. This parallels commonly observed heterogeneity across iPSCs in gene expression and DNA methylation, as well as phenotypes such as growth rate and cellular morphology (Volpato & Webber, 2020). There have been many efforts to evaluate the quality of individual iPSC lines. In mouse iPSCs, chimera formation can be used to determine the ability of cells to form adult tissue. Ultimately, the most stringent test is tetraploid complementation, where cells must lead to development of a whole organism (Robinton & Daley, 2012). Complete reprogramming to reconstitute an entire organism has been shown through the cloning of numerous animals in different species (Gurdon & Byrne, 2004), consistent with the high developmental potential of human NT-ESCs. In contrast, only a few studies reported full developmental competence of mouse iPSC lines reprogrammed from adult cells in tetraploid embryo complementation assays (Stadtfeld et al., 2012). Whether these differences between nuclear transfer ES cells and iPSCs are due to a selection for developmental potential during early development, or due to other mechanisms, is not known. Specific challenges arise in the human system, as stringent developmental assays are not available for human iPSCs. Instead, criteria for selection of human iPSCs must rely on *in vitro* differentiation potential and detailed molecular characterization. However, assays for gene expression and DNA methylation largely fail to separate ESCs and iPSCs (Bock et al., 2011; Johannesson et al., 2014; Koyanagi-Aoi et al., 2013; Kyttälä et al., 2016), suggesting there is not an iPSC-specific signature of these properties. While some genes and DMRs can serve as markers for reduced iPSC differentiation potential, they represent a small number of loci (Koyanagi-Aoi et al., 2013). Moreover, even signatures of epigenetic memory for iPSCs derived from a given cell type are inconsistent across studies (Robinton & Daley, 2012), leaving a need for a more consistent and readily attainable molecular signature. To integrate multiple molecular measurements, the Pluritest (Müller et al., 2011) and a “score card” for deviation from ES cells (Bock et al., 2011) bioinformatic tools have been developed, however they only measure pluripotency and not differentiation capacity (Liu & Zheng, 2019) and rely on labor- and cost-intensive data collection that may not be easily scalable. Furthermore, these methods often disagree with one another (Allison et al., 2018; Bouma et al., 2017).

While further work is needed to examine the links between replication timing and stem cell pluripotency, we suggest that replication timing could potentially be used as a screen for iPSC quality in parallel to other quality controls and without the obligatory need for RNA-seq or bisulfite analysis. Our results suggest that reproducible replication timing aberrations are broader than gene expression and DNA methylation signatures in iPSCs. Furthermore, DNA replication timing profiling can be performed using the same data (WGS) used for genotyping and identification of *de novo* mutations, making it a compelling molecular signature at minimal added costs for identifying low quality iPSCs prior to use in research or cell therapy.

## Methods

### Stem cell derivation and culture

All experiments with human pluripotent stem cells were reviewed and approved by the Columbia Institutional Review Board (IRB) and the Columbia University Embryonic Stem Cell Committee. Human pluripotent stem cell lines were cultured on geltrex-coated plates in StemFlex Medium (A3349401, Gibco) to 80-90% confluence and passaged with TrypLE (12605036; Life Technologies). iPS cell lines were generated using mRNA mediated reprogramming (Johannesson et al., 2014). The cell line 1000-5 (which was excluded from the analysis) was generated using Sendai viral vectors. Previous studies had not observed significant differences between iPSCs generated using different non-integrating reprogramming methods (Schlaeger et al., 2015).

### NPC differentiation

Cells were differentiated to NPCs from human pluripotent stem cell lines using a published protocol with modifications (Wang et al., 2016). Briefly, human pluripotent stem cells were cultured in 6-well geltrex-coated dishes with StemFlex medium. When each well reached 95-100% confluence, StemFlex medium was replaced with KSR differentiation medium; this was considered day 0 of differentiation. The KSR differentiation medium was changed daily for 4 days. From day 5 to 7, cells were incubated with KSR differentiation medium mixed with N2 differentiation medium at ratios of 3:1 (day 5), 1:1 (day 6), and 1:3 (day 7). On day 8, detection of EdU staining and PAX6 expression by FACS was performed (**Fig. S1C**), and cell pellets were collected for whole genome sequencing.

KSR differentiation medium consisted of knockout DMEM (10829-018, Gibco) supplemented with 15% KSR (10828-028, Gibco), 1% GlutaMAX (35050061, Thermo Fisher), 1% NEAA (11140-050, Gibco), 0.1 mM 2-Mercaptoethanol (21985-023, Gibco), 10,000 U/ml penicillin–streptomycin (15070-063, Thermo Fisher), 10 μM SB 431542 (S1067, Selleckchem), and 2.5 μM LDN 193189 (S2618; Selleckchem). N2 differentiation medium was based on DMEM/F12 (10565042, Thermo Fisher) by supplementing with 1% N2 supplement (17502-048, Thermo Fisher), 1% GlutaMAX, 1% NEAA, 0.4 N ascorbic acid (A-5960, Sigma), 1% D-glucose with 16% (w/v) (G8270, Sigma) 10,000 U/ml penicillin–streptomycin, 10 μM SB431542 (S1067, Selleckchem), and 2.5 μM LDN193189 (S2618; Selleckchem).

### FACS

EdU staining was performed by Click-iT^TM^ EdU Alexa Fluor 488 Imaging Kit (C10337), samples were washed with cold PBS and centrifuged for 5 min at 1000 rpm at 4 °C. Samples were incubated with PAX6 antibody (901302, Biolegend; **Fig. S1C**) at room temperature (RT) for 30 min, followed by 3 washes with PBST (3% BSA in PBS with 0.1% Triton) and a secondary Alexa Fluor 647 donkey anti-rabbit IgG (H+L) (A31573, Thermo Fisher) for another 30 min at RT. The cell suspension was analyzed on a BD Fortessa flow cytometer, and data was analyzed using FlowJo v.10.

### Whole genome sequencing

Samples were sequenced in two batches: batch 1 included 26 of the 28 ESCs and 17 of the 19 iPSCs, and batch 2 included the remaining two ESCs and two iPSCs, and all 17 NPCs. For both batches, DNA was extracted with the MasterPure DNA purification kit (Lucigen). Libraries were prepared using the Illumina TruSeq PCR-free library preparation kit, and sequencing was performed on the Illumina HiSeq X Ten with 150-bp paired-end reads at GeneWiz (South Plainfield, NJ). Reads were then aligned to the hg19 human reference genome using BWA-MEM (Li & Durbin, 2010).

### Replication timing profiles

DNA replication timing was inferred as described in (Edwards et al., 2021). Genome STRiP (Handsaker et al., 2015; Koren et al., 2014) was used to infer DNA replication timing across the genome. Sequence read depth was calculated in 2.5-kb windows along the genome, corrected for alignability and GC content. Copy number values were normalized to an average DNA copy number of two. These copy number values were then filtered as follows:

1. Windows spanning gaps in the reference genome were removed.
2. Windows with copy number greater than one above or below the median copy number were removed.
3. In order to remove extreme data points, the data was segmented using the MATLAB function *segment*, which groups consecutive data points into segments based on a tolerance threshold. This analysis was done twice using two different segmentations parameters of 0.5 (less strict) and 0.1 (more strict). By using two different parameters, both shorter and larger genomic regions that deviate from the median can be captured. Segments falling above or below the median by a threshold of 0.45 copies were removed.
4. Genomic regions that were further than 30 kb from other data points and that were <300 kb long were removed.
5. Regions shorter than 100 kb between removed data points were removed.
6. Regions shorter than 500 kb between three or more removed data points were removed.

Data were then smoothed using the MATLAB function *csaps* with smoothing parameter of 10^−17^, and then normalized to an autosomal median of 0 and a standard deviation of 1, such that positive values represent early replication and negative values represent late replication. After the above filtering steps, the iPSC cell line BJiPSM still showed a reduced copy number on a portion of the p arm of chromosome 18 (chr18:1-10063413) and an elevated copy number on the entire q arm of chromosome 9 (chr9: 70938816-141135844); these regions are suspected to be subclonal aneuploidies and were removed from further analysis.

The mean correlation of replication timing in ESC and iPSC samples from the second batch (with the exception of cell line 1000-5 that was removed due to low correlations with other cell lines and being an outlier by PCA) with the first batch was r=0.96, equal to the internal correlation of batch 1 (r=0.96); this suggests little if any batch effects. PCA analysis also clustered all ESC and iPSC samples together, as well as NPCs together (PC1 = 89.1% explained, PC2 = 5.7% explained), with the only clear visual outliers being the removed samples: 1000-5, as well as the three NPCs labeled in **Fig. S1A**.

### Replication timing variation identified using filtering-based ANOVA-scan

To call replication timing variation between NT-ESCs and isogenic iPSCs, we used a modified version of the ANOVA-based scan with filtering that we used previously (Edwards et al., 2021). Briefly, regions of 76 replication timing windows (covering 190kb of uniquely alignable sequence) were tested, with a slide of a quarter region (such that each genomic locus was tested four times) along the entire genome using a one-way ANOVA test between the relevant groups (e.g., NT-ESCs versus isogenic iPSCs). A Bonferroni corrected p-value cutoff of p < 9.58 × 10^−7^ (0.05/52,184 regions tested) was employed, and regions that were overlapping or separated by short (< 750kb) gaps were merged. Because there were more samples than in our previous work (Edwards et al., 2021), the filtering step was modified to require that pairwise comparisons of NT-ESC and iPSC profiles showed a consistent direction of effect (delayed or advanced relative to the mean of the other group) in at least 3 samples (75%) for each group (i.e., at least 3 NT-ESC were required to be earlier/later than the mean of iPSCs, and conversely iPSCs were reciprocally later/earlier with respect to the NT-ESC mean). As with the original scan, windows where profiles overlapped were removed, and only those regions greater than 76 windows in size were maintained. Finally, to capture the full region of variation, all variant regions were extended bidirectionally as long as the two sample groups remained separated and the ANOVA p-value was still below the significance threshold. This resulted in the final set of 24 NT-ESC/iPSC variants and identified no variants in the ESC/NT-ESC comparison.

### Down-sampling analysis for filtering-based ANOVA-scan

There were 24 replication timing variants between the four NT-ESCs and four isogenic iPSCs, while a comparison of the four NT-ESCs with all 28 ESCs (using the more appropriate empirical scan for larger samples) did not identify any variation. To test that this was not a function of the number of cell lines, or the scan used, the following down-sampling analysis was performed:

Four randomly selected samples from each stem cell group (ESC, NT-ESC, iPSC) were compared using the initial filtering-based ANOVA scan. This was performed 100 times for each comparison, and the mean variation for each comparison was calculated. The mean extent of variation called between NT-ESCs/down-sampled iPSCs (0.79% autosomal span) was identical to that identified between NT-ESCs/isogenic iPSCs (0.79% autosomal span; **Fig. 1B**), indicating that the full set of iPSCs exhibited similar variation to the isogenic subset. Down-sampled ESC/NT-ESC mean variation was lower (0.4%), consistent with the lack of variation seen when comparing the full set of ESCs to NT-ESCs. Additionally, intra-cell-type comparisons of ESCs (4 ESCs v 4 ESCs) and iPSCs (4 iPSC vs 4 iPSC) both revealed 0.38% variation, indicating that ESC/NT-ESC variation is akin to comparisons between cells of the same type. Finally, the mean down-sampled ESC/iPSC variation was greater than all other down-sampled comparisons (1.2%), consistent with the greater variation we identified between these samples. Thus, regardless of the scan methodology used (the filtering approached used here, or the empirical approach used to identify ESC/iPSC variants) and the sample size, we still observe minimal ESC/NT-ESC variation, and notable iPSC/ESC variation.

### Replication timing variation identified using empirical ANOVA-scan

In order to identify replication timing variation using a larger number of samples (ESC/NT-ESC and ESC/iPSC comparisons), a permutation-based approach was employed using the genome-wide ANOVA scan to establish an empirical p-value cutoff. Because of the heterogeneity of the iPSC samples, permuting ESC and iPSC samples together would result in high levels of background variation, and therefore only ESC samples were permuted to quantify the expected level of variation in cell lines without cell type differences. One thousand permutations of the 28 ESC samples were performed, comparing a random selection of 14 samples to the remaining 14 in each permutation using the ANOVA scan (see above description of filtering based scan, and (Edwards et al., 2021) for a detailed description; the filtering steps for direction of effect and profile overlap were excluded from this empirical analysis). One additional modification was made, wherein initial windows were tested with the one-way ANOVA, and then corrected with a q-value approach (Storey, 2002). Adjacent windows with q-values <0.05 were merged, and then still subjected to a one-way ANOVA which was required to pass the genome-wide Bonferroni-corrected threshold. By using this multistep approach, large regions with subtle variation that may have escape the initial cutoff in 190kb windows, but which may be significant, can be identified.

The final p-value cutoff was determined by an empirically determined threshold at a False discovery rate (FDR) < 0.05. FDR was calculated by dividing the total variation found (in Mb) by the average variation across 1000 permutations. This was then performed at every p-value cutoff (lowering by 10^-1^ each time) until the false discovery rate of 0.05 was reached at p = 1 x 10^-53^. Regions in the true comparison under this threshold were then extended in both directions until profiles overlapped or the p-value dropped below the threshold to form the final variant list.

To ensure that this estimate of false variation was not influenced by the sample size differences (14 vs 14 in permutations compared to 18 vs 28 in ESC/iPSC comparisons), a permutation analysis was also performed on the 108 validation ESCs. Nine of the 108 ESCs were previously shown to have autosomal replication delays linked to putative X reactivation (Edwards et al., 2021) and were excluded from this analysis. A hundred permutations were performed comparing 14 vs 14 of these ESCs, and separately 100 permutations comparing 18 and 28 ESCs. At a cutoff of p = 1 x 10^-53^, the 18 vs 28 comparison resulted in an FDR of 0.054, while the 14 vs 14 comparison resulted in FDR = 0.024. Thus, while the larger sample size may result in slightly more background variation, false discovery rates near 0.05 are nonetheless observed at this p-value cutoff even with larger numbers of samples.

NT-ESCs were also compared to the full set of 17 iPSCs using the empirical approach. While this did not produce any variation at 5% FDR (likely due to a high degree of false positives resulting from iPSC variability, see **Fig. 2**), the full set of iPSCs directly validated the variants found using the isogenic samples (**Fig. S2**).

### ESC/iPSC variant validation

Replication timing profiles (corrected using principal component analysis) for the 108 ESCs and 300 iPSCs were obtained from Ding et al. 2021. Because ESC profiles were generated using 10kb replication timing windows, while iPSC profiles were generated using 2.5kb profiles, the iPSC profiles were interpolated to the ESC coordinated prior to analysis. In order to test whether iPSCs exhibited variation in these regions compared to ESCs in the validation set, we first ensured that these regions were consistent between datasets by calculating the mean Pearson correlations between the validation ESCs and the original ESCs for each variant and all non-variant regions. In non-variant regions the average correlation was r = 0.83. Testable regions were defined as those where ESC/ESC correlation was r > 0.70, leaving 15 of the 22 iPSC-delayed and two of four iPSC-advanced variants for testing.

Twelve of the 15 iPSC-delayed variants and both iPSC-advanced variants tested showed the expected direction of effect (delayed/advanced) across 50% or more of the variant region. Suspected batch effects precluded testing for significant variation using ANOVA. However, these regions were tested by asking whether replication timing difference between the iPSC mean and ESC mean in each region was more extreme than flanking regions of equal size to the variant. Flanking regions were required to contain at least half as many data points as the variant region, i.e., be less than 50% spanned by a reference genome gap. Variants where the maximum replication timing difference (iPSC vs. ESC) was more extreme (more positive for advanced regions or more negative for delayed regions) were considered validated.

### Permutation methodology

In order to determine the significance of overlap between replication timing variants and various other genomic features, replication timing variant locations were permuted 1000 times, and significance was determined by comparing the overlap between variants and the tested feature of interest to permutations.

For generating permuted variant regions, we required the following:

1. Each permutation consisted of a number of permuted windows equal to the number of variants, and each permuted window was the same size as the variant from which it was derived.
2. Replication timing in the middle of the permuted windows was required to be within ± 0.2 SD of the variant from which it was derived.
3. Permuted regions could not overlap variants or each other. Permuted regions were ordered by the p-value of the corresponding variant, so the most significant variant was permuted first, and later regions within this same permutation could not overlap the regions that were determined prior.
4. Permuted regions could not overlap gaps in the reference genome.
5. Regions had to have data in at least 50% of the replication timing bins.

Overlap was determined in one of two ways. For comparisons with features of large (Mb-scale) size (comparisons of ESC/iPSC variants to NT-ESC/iPSC variants, non-GC DMRs, and NPC variants) the overlap was considered as a cumulative percentage of each variant. For example, if 40% of a replication timing variant coincided with a region of interest, the contribution of this region to the total overlap would be 0.4. Summed over all the variants, this meant that the total possible overlap for *n* variants ranged from 0 to *n*. Doing so normalized the contribution of each variant equally, regardless of size. Overlap comparison for smaller (sub Mb scale) features (those between ESC/iPSC variants and CG-DMRs, genes, or ChrommHMM states which were considered as a series of single base pair states) were simply a cumulative count of the number of features which fell inside variant regions.

### Fibroblast replication timing profiles

Using single cell DNA sequencing data of BJ fibroblast (isogenic to NT5/6/8, BJiPSM, and BJiPSO; 10xgenomics.com), we identified cells in S phase and aggregated their DNA sequence data to generate a profile of fibroblast replication timing as previously described (Massey & Koren, 2022). Briefly, G1/G2 cells and S phase cells were identified based on the fluctuations in read depth along chromosomes in the sequence data. Genomic windows were constructed using the G1/G2 cells, and for each window, each S phase cell was categorized as either having replicated or non-replicated DNA. Aggregating the single cell profiles created an ensemble profile that was used for analysis.

Accordingly, the single-cell fibroblast profile was used for comparison with ESC and iPSC replication timing. Six variants were tested that had a correlation of r > 0.3 between the fibroblast and ESC and iPSC means, and where the ESC or iPSC mean showed replication timing similar to fibroblasts (+/-0.2 SD) at the region of maximum variation between ESCs and iPSCs. These variants were then categorized based on whether the difference in replication timing was smaller between the iPSC mean and fibroblast (epigenetic memory) or between the ESC mean and fibroblast (reprogramming-linked aberration). In all cases, fibroblasts were only within 0.2 SD of either ESCs or iPSCs, but not both.

### DNA methylation data and DMRs

All CG DMRs (Lister et al., 2011) were successfully converted from hg18 to hg19 using the UCSC LiftOver tool, while nineteen of 22 non-CG iPSC hypomethylated, and six of seven non-CG iPSC hypermethylated regions were successfully converted to hg19. In order to obtain raw methylation data for plotting purposes (**Fig. 4C**), the 26 variant regions as well as 1Mb upstream and downstream were first lifted from hg19 to hg18. Four of these regions failed to be lifted over, leaving 22 variant regions for analysis. Methylation data in these regions was then obtained and lifted back to hg19. All data in **Fig. 4C** was plotted using hg19 coordinates. A flanking region size of 1Mb was chosen (instead of 3Mb as in the remainder of our analyses) because larger flanking regions failed LiftOver.

Non-CG methylation data was processed according to methods described previously(Lister et al., 2011), differing only in the size of the flanking region. Each variant region +/- 1Mb was divided into 10 Kb windows, and the methylation level of all CHH and CHG context sites was calculated. These window values were then normalized to the maximum value across the region, and data was then averaged for ESCs and iPSC before plotting.

## Acknowledgments

This work was supported by NYSTEM grant #C32564GG and the US Israel Binational Science Foundation grant #2015089 to DE, and in part by the National Institutes of Health grant R35GM148071 to AK. NW was supported by a P.R. China Scholarship Council fellowship. MME was supported as a Cornell Center for Vertebrate Genomics distinguished scholar.

## Data availability

All raw sequencing data generated in this study have been submitted to the NCBI Database of Genotypes and Phenotypes (dbGaP; https://www.ncbi.nlm.nih.gov/gap/) under accession number phs001957.

