## Supplemental Figures for "Incomplete Reprogramming of DNA Replication Timing in Induced Pluripotent Stem Cells"

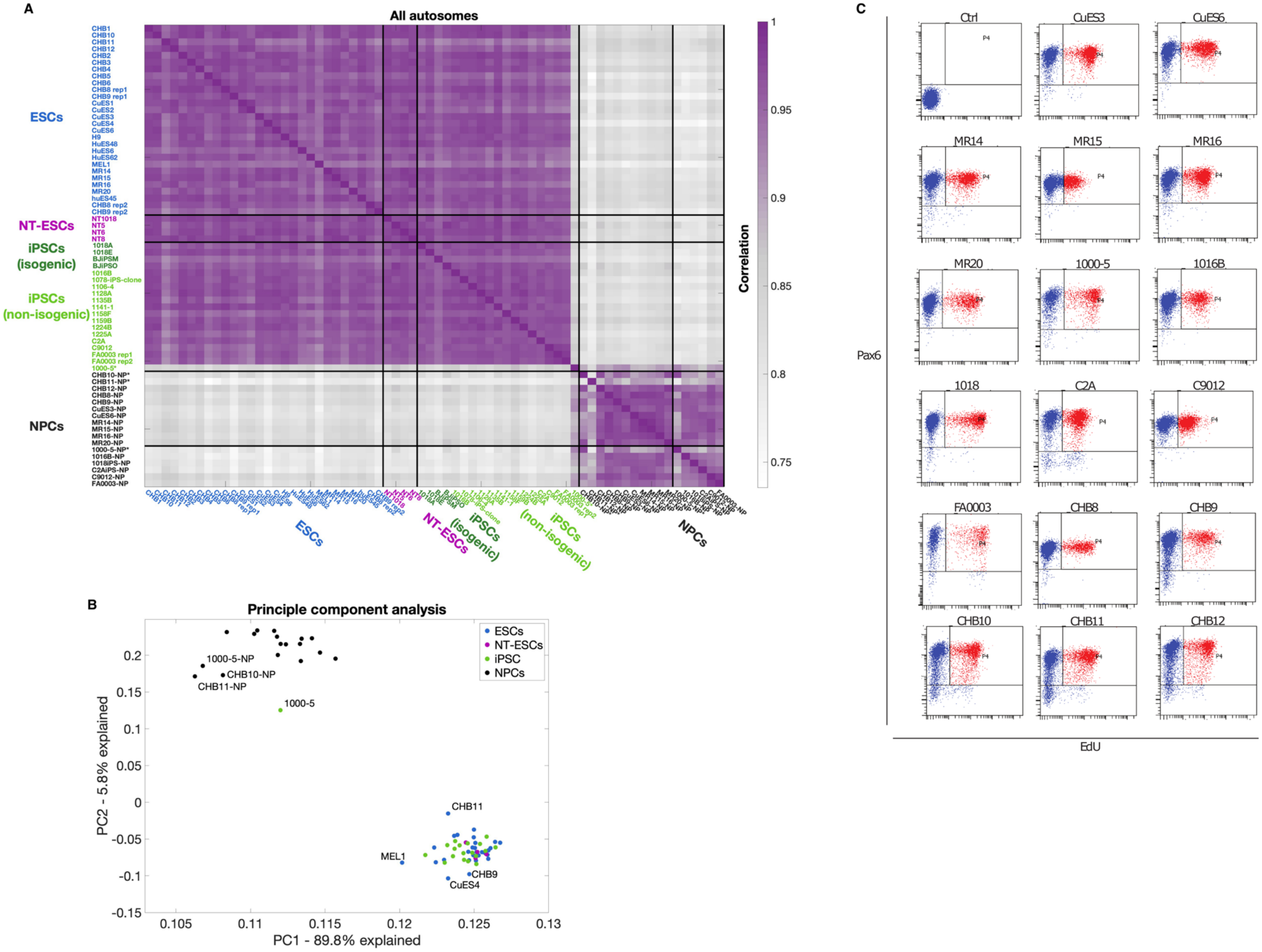

A
All autosomes
ESCs
NT-ESCs
iPSCs (isogenic)
iPSCs (non-isogenic)
NPCs
Correlation
1
0.95
0.9
0.85
0.8
0.75
B
Principle component analysis
ESCs
NT-ESCs
iPSC
NPCs
1000-5-NP
CHB10-NP
CHB11-NP
1000-5
CHB11
MEL1
CHB9
CuES4
PC2 - 5.8% explained
PC1 - 89.8% explained
C
Ctrl
CuES3
CuES6
MR14
MR15
MR16
MR20
1000-5
1016B
1018
C2A
C9012
FA0003
CHB8
CHB9
CHB10
CHB11
CHB12
Pax6
EdU

**Supplementary Figure 1: Replication Timing Profile Quality Assessment. A.** Correlation matrix of DNA replication timing for 28 ESCs (blue), four NT-ESCs (purple), four iPSCs isogenic to NT-ESCs (dark green), 15 additional iPSCs (green), and 17 NPCs (black) across the autosomes. **B.** Principal component analysis (PCA) of autosome-wide replication timing for all samples. Stem cells (ESC: blue, iPSC: green, NT-ESC: purple) cluster separately from NPCs (black). Cell lines marked with * were removed from further analysis: The iPSC line 1000-5 did not cluster with other stem cells, instead showing greater resemblance to NPCs, consistent with its higher correlation to NPCs in A; This cell line is suspected to have spontaneously differentiated; The NPC lines 1000-5-NP, CHB10-NP, and CHD11-NP, clustered separately from, and showed reduced correlation to, other NPCs (r = 0.91, 0.92, and 0.88 to all NPCs, respectively, compared to an average of r = 0.96 for all other NPCs). C Flow cytometry analysis for indicated neuronal progenitor cells marker Pax6 from day 8 differentiation, as well as a marker EdU of DNA synthesis during the S-phase of the cell cycle. The Pax6 and EdU percentages of neuronal progenitor cells in **Supplementary Table 1**.

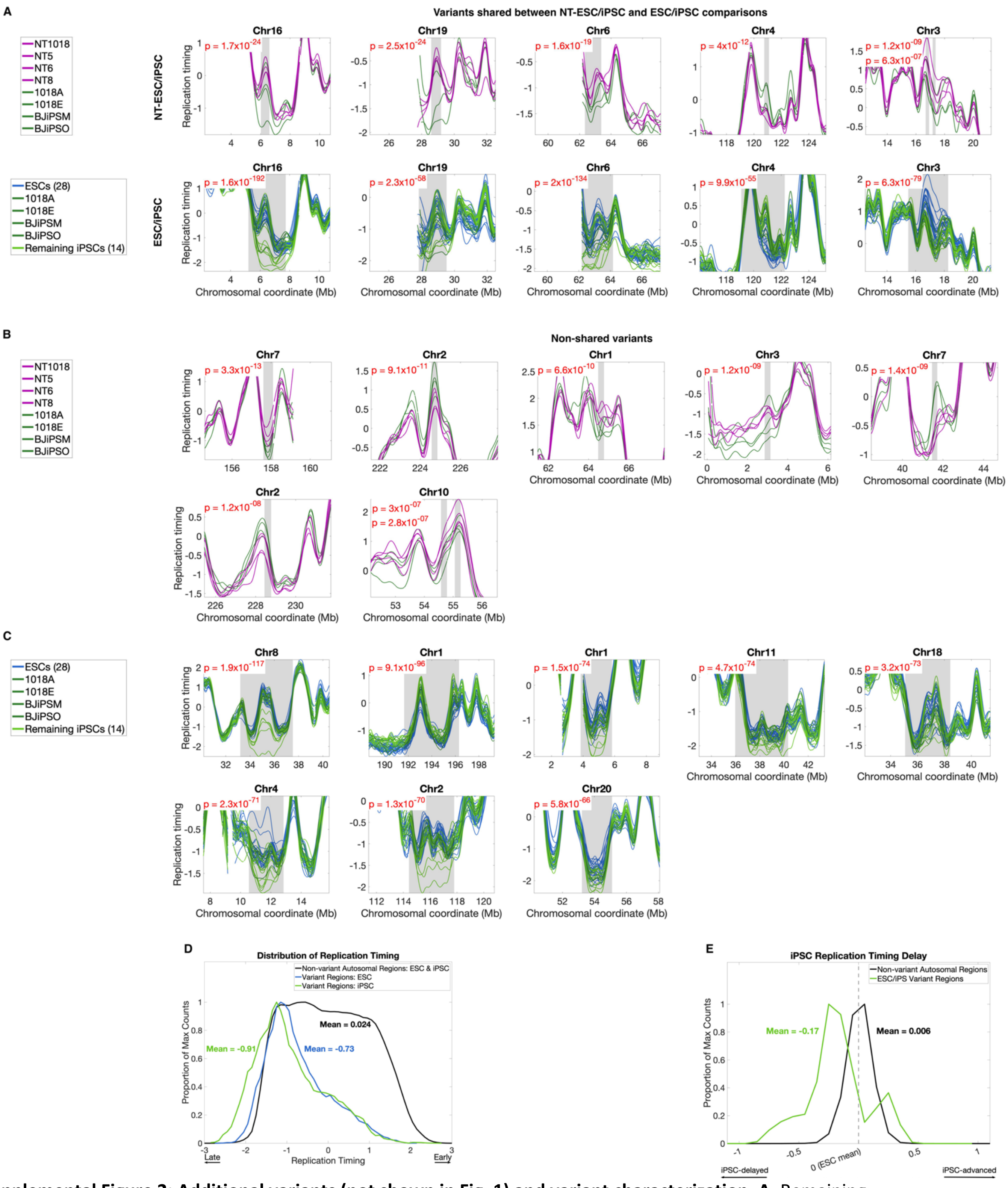


**Supplemental Figure 2: Additional variants (not shown in Fig. 1) and variant characterization. A.** Remaining overlapping NT-ESCs/iPSC and ESC/iPSC variants not shown in **Fig. 1B-C.** Six NT-ESC/iPSC variants overlap five ESC/iPSC variants. Two NT-ESC/IPSC variants on chromosome 3 are shown together due to proximity, and p-values in these two plots are ordered with the leftmost variant's p-value listed first. **B.** Remaining NT-ESC/iPSC variants which do not overlap ESC/iPSC variants. Two NT-ESC/IPSC variants on chromosome 10 are shown together due to proximity, and p-values in these two plots are ordered with the leftmost variant's p-value listed first. **C.** Remaining ESC/iPSC variants that do not overlap NT-ESC/iPSC variants. **D.** Distribution of the replication timing difference (mean across all samples) between ESCs and iPSCs at variant regions (green) and non-variant regions (black). Histograms are normalized to the maximum values. **E.** Distribution of replication timing in ESCs (blue) and iPSCs (green) at variant regions, and of both ESCs and iPSCs in non-variant regions (black). Histograms are normalized to the maximum values**.**

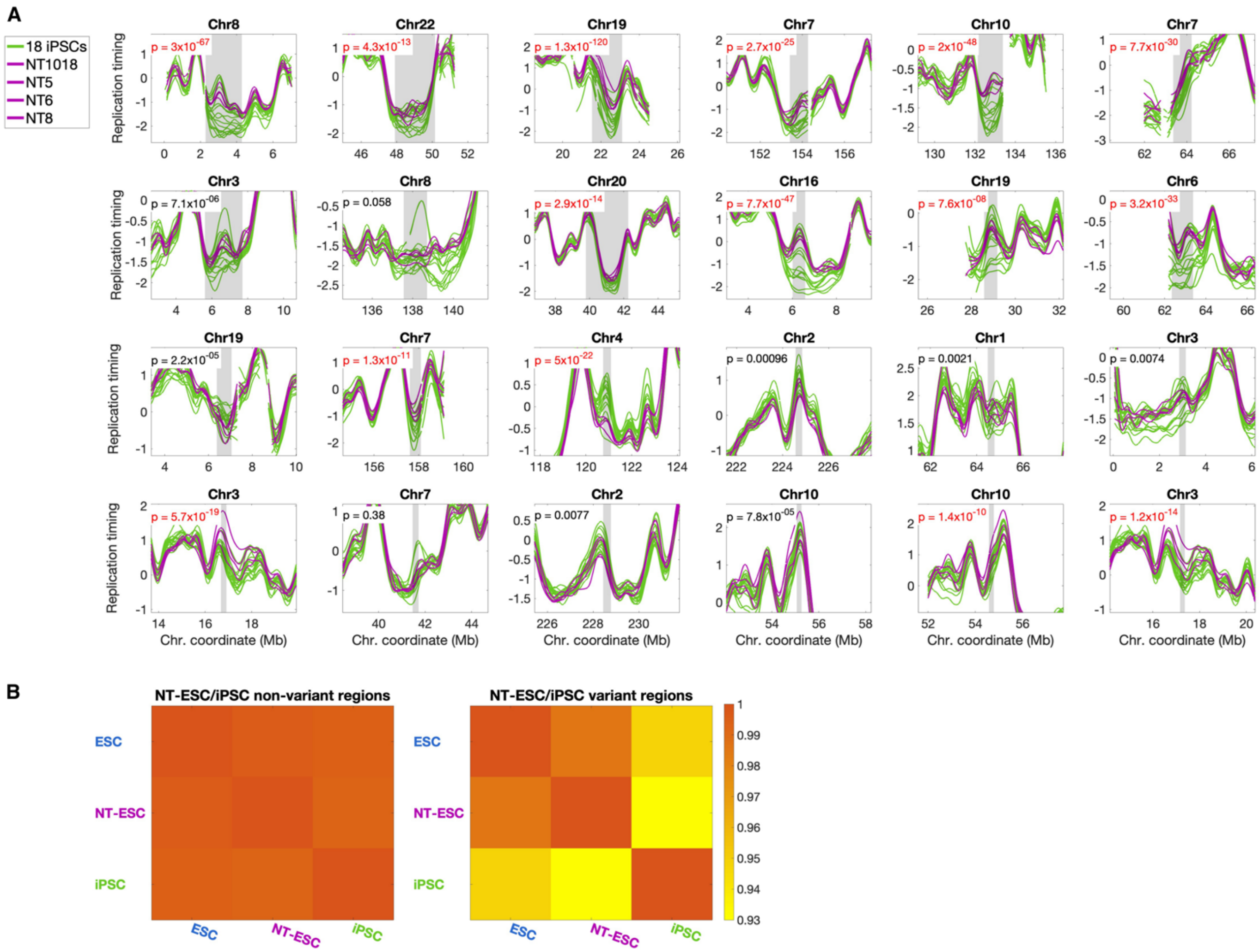


**Supplemental Figure 3: Validation of NT-ESC vs iPSC variation using all 18 iPSCs**. **A**. Direct testing of variation between NT-ESCs and 18 iPSCs at all 24 NT-ESC/iPSC variants initially identified using only the four iPSC cell lines isogenic to the NT-ESCs. Variants are ordered by initial NT-ESC/iPSC p-value. Each variant region was tested using ANOVA, and p-values are reported in the top left corner (significant p-values less than the genome-wide Bonferroni corrected threshold of 9.13 x $10^{-7}$ were observed for 15/24 of the variants; labeled in red). **B.** The mean correlations of replication timing profiles for the three stem cell types at autosomal regions outside NT-ESC/iPSC variants (left) and at NT-ESC/iPSC variants regions (right). While replication timing is highly consistent at non-variant regions, the mean of all 18 iPSCs shows significantly lower correlation to both ESCs and NT-ESCs at variant regions.

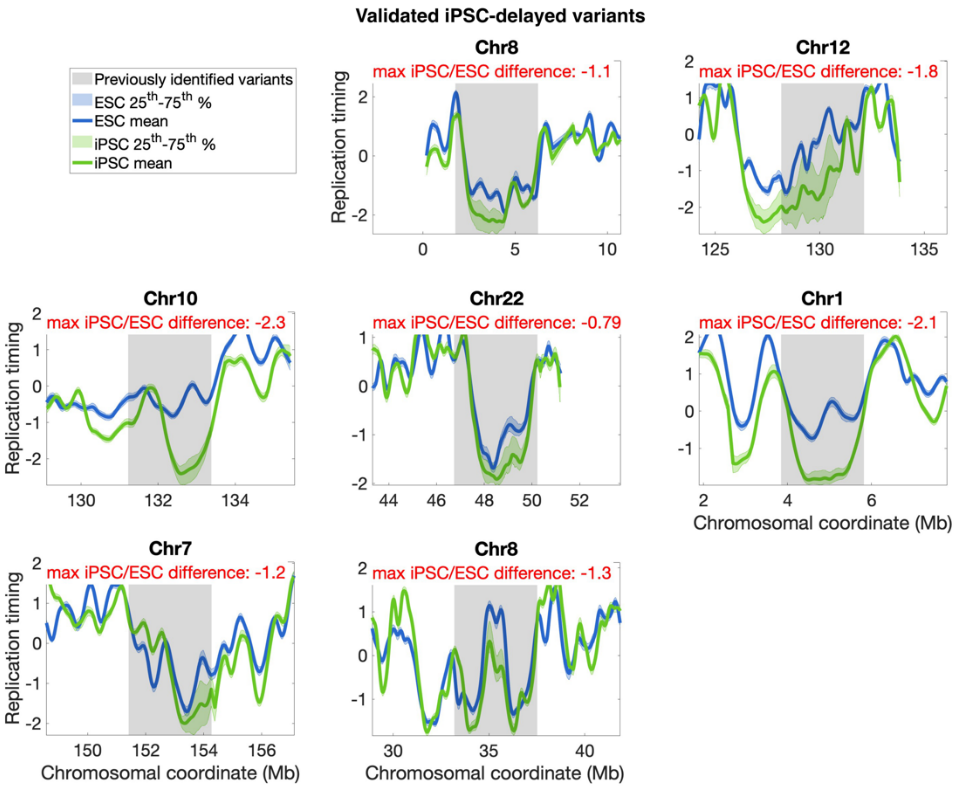


**Supplemental Figure 4: Replication timing variation between ESCs and iPSCs in a larger cohort of stem cell lines**. Replication timing profiles of 108 ESCs and 300 iPSC at the seven validated variant regions. The magnitude of replication timing differences between the stem cell types is indicated at top (standard deviation units). The 25th to 75th percentile are shown alongside the mean replication timing profiles for each stem cell type.

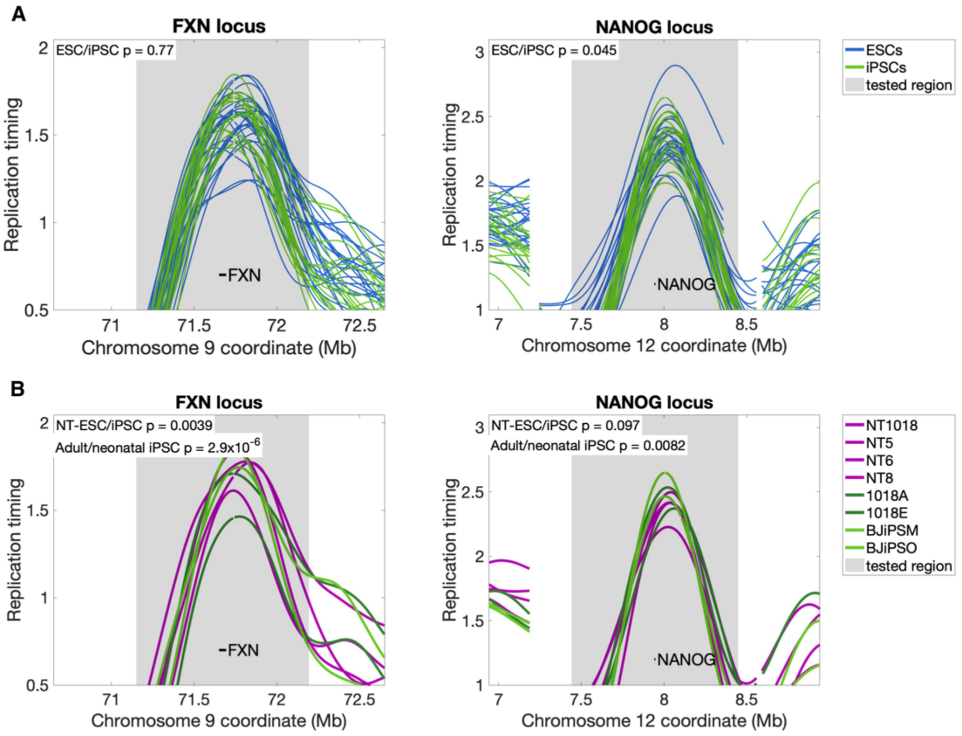


**Supplemental figure 5: Several known differentially expressed genes are found in variants regions, while FXN and NANOG loci show little to no replication timing variation between ESCs and iPSCs**. **A.** Replication timing profiles for ESCs and iPSCs at the FXN and NANOG loci. The region +/- 500Kb from each gene was tested for variation using ANOVA. While ESC/iPSC variation was significant at a 0.05 threshold for the NANOG locus, neither locus showed significant variation at the genome-wide Bonferroni-corrected threshold. **B**. Same as above for NT-ESCs and isogenic iPSCs (the same ones studied by Paniza et al., 2020). NT-ESC/iPSC differences at both loci were significant at a 0.05 threshold, though neither showed significant variation at the genome-wide Bonferroni-corrected threshold. We also considered adult/neonatal iPSC differences, which were significant at a 0.05 threshold for the FXN locus but were not significant for either locus at a genome-wide Bonferroni corrected threshold.

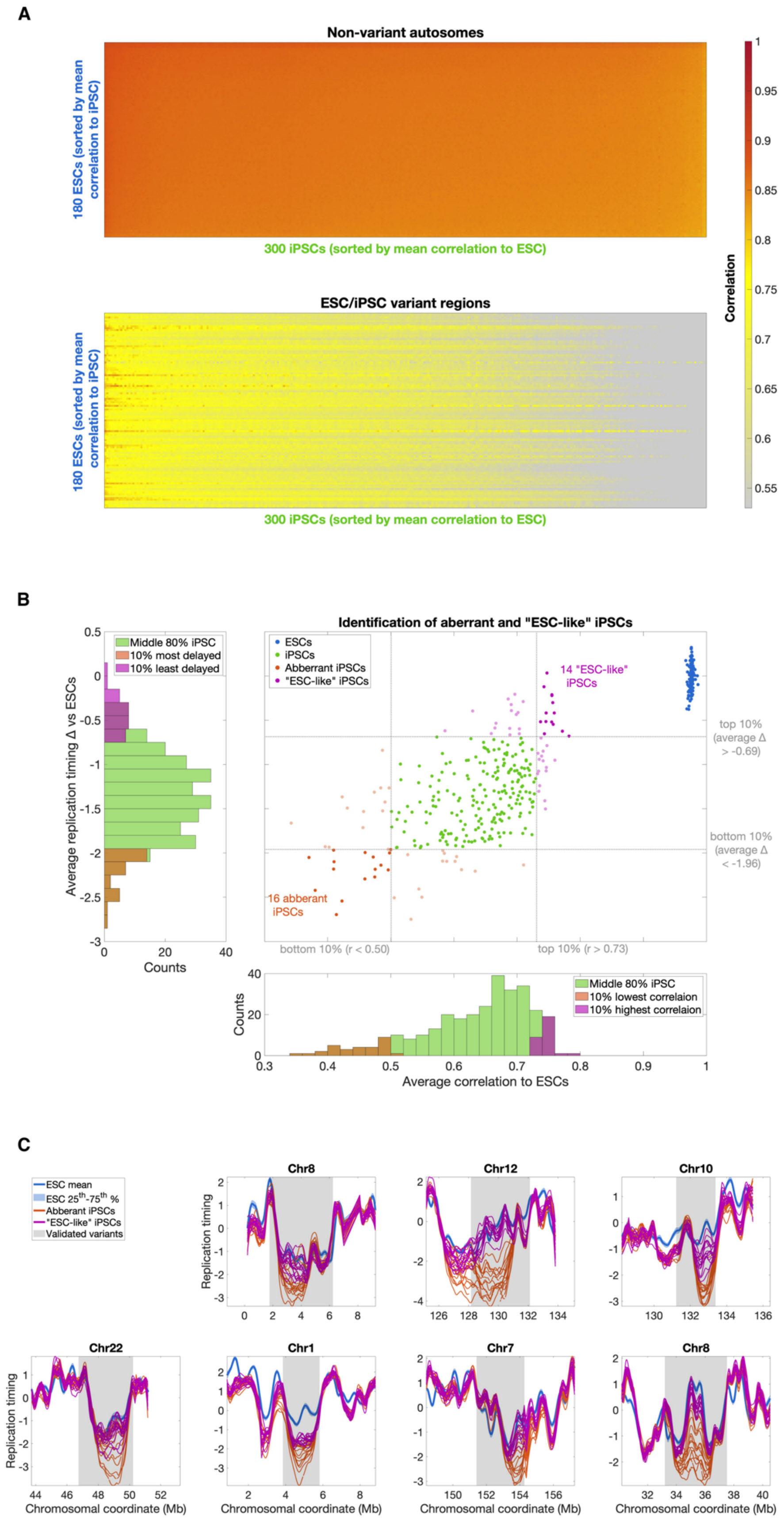
A
Non-variant autosomes
180 ESCs (sorted by mean correlation to iPSC)
300 iPSCs (sorted by mean correlation to ESC)
ESC/iPSC variant regions
180 ESCs (sorted by mean correlation to iPSC)
300 iPSCs (sorted by mean correlation to ESC)
Correlation
1
0.95
0.9
0.85
0.8
0.75
0.7
0.65
0.6
0.55
B
Identification of aberrant and "ESC-like" iPSCs
Middle 80% iPSC
10% most delayed
10% least delayed
Average replication timing Δ vs ESCs
Counts
ESCs
iPSCs
Abberrant iPSCs
"ESC-like" iPSCs
14 "ESC-like" iPSCs
16 abberant iPSCs
top 10% (average Δ > -0.69)
bottom 10% (average Δ < -1.96)
bottom 10% (r < 0.50)
top 10% (r > 0.73)
Middle 80% iPSC
10% lowest correlaion
10% highest correlaion
Average correlation to ESCs
C
ESC mean
ESC 25th-75th %
Abberant iPSCs
"ESC-like" iPSCs
Validated variants
Chr8
Chr12
Chr10
Chr22
Chr1
Chr7
Chr8
Replication timing
Chromosomal coordinate (Mb)

**Supplemental Figure 6. Identification of aberrantly replicating and ESC-like iPSCs among the larger cohort of stem cell lines. A.** Correlation of replication timing between 108 ESCs (rows) and 300 iPSCs (columns) at non-variant regions (top) and validated ESC/iPSC variant regions (bottom). **B.** Definition of aberrant and "ESC-like" iPSC lines based on correlations to ESC replication timing profiles (bottom histogram; x-axis) and average replication difference from the ESC mean (labeled as Δ) at validated ESC/iPSC variant regions (left histogram; y-axis). In the histograms and the scatter plot, the top 10%, bottom 10%, and remainder iPSCs are shown in orange, purple, and green, respectively. Inclusion in both top or both bottom deciles was used for defining the ESC-like and aberrant iPSC categories. **C.** Aberrant (orange) and ESC-like iPSCs (purple) compared to the inter-quartile range of ESCs (blue) at all ESC/iPSC replication timing variants validated using this dataset (**Fig. S4A)**.

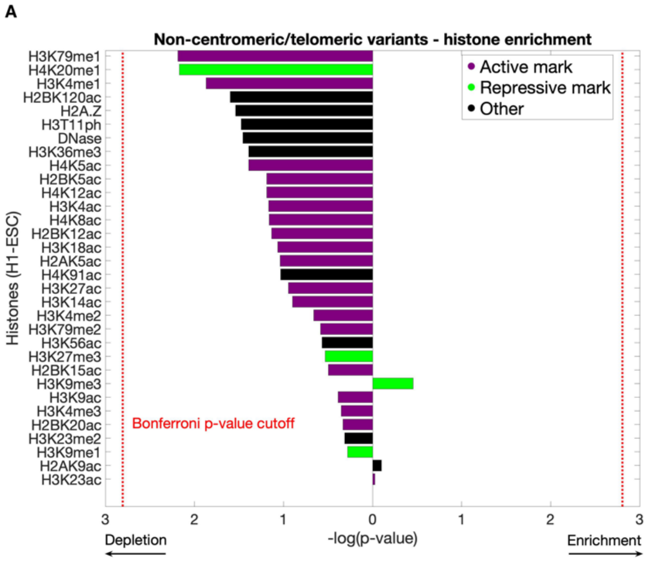


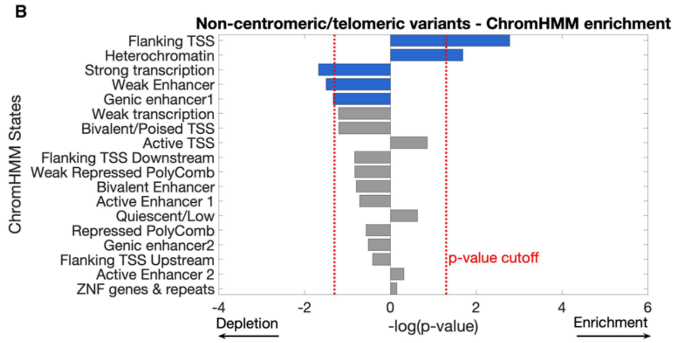


**Supplemental Figure 7: Differentially expressed genes and additional heterochromatin enrichment in variant regions. A.** Histone enrichment in non-centromeric/telomeric variants, similar to **Fig. 3C. B.** ChromHMM enrichment in non-centromeric/telomeric variants, similar to **Fig. 3D**.

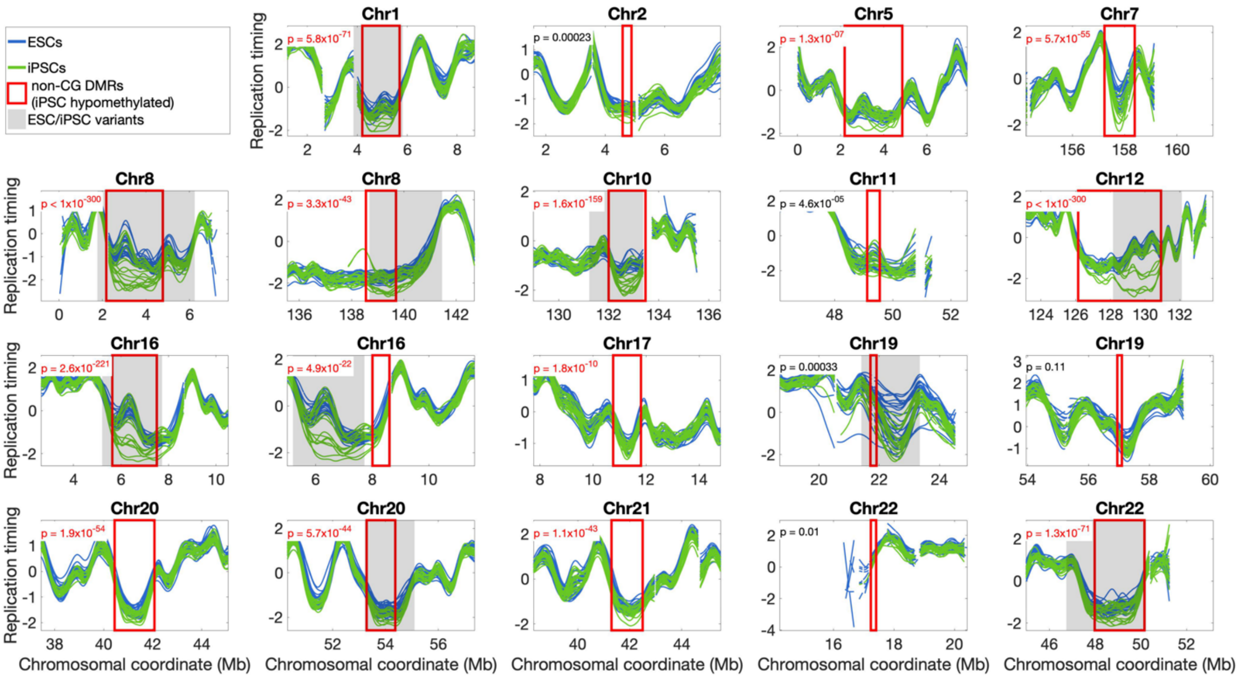


**Supplemental Figure 8 Replication timing in ESCs/iPSCs at non-CG DMRs:** Direct testing of ESC/iPSC replication timing variation at non-CG DMRs where iPSCs are hypomethylated (red). Grey: ESC/iPSC replication timing variants. Significant p-values at a Bonferroni corrected genome-wide threshold are shown in red, while non-significant p-values are in black. All eight DMRs overlapping variant regions are significant, as expected, as are an additional six which were not identified in the original replication timing variant scan.

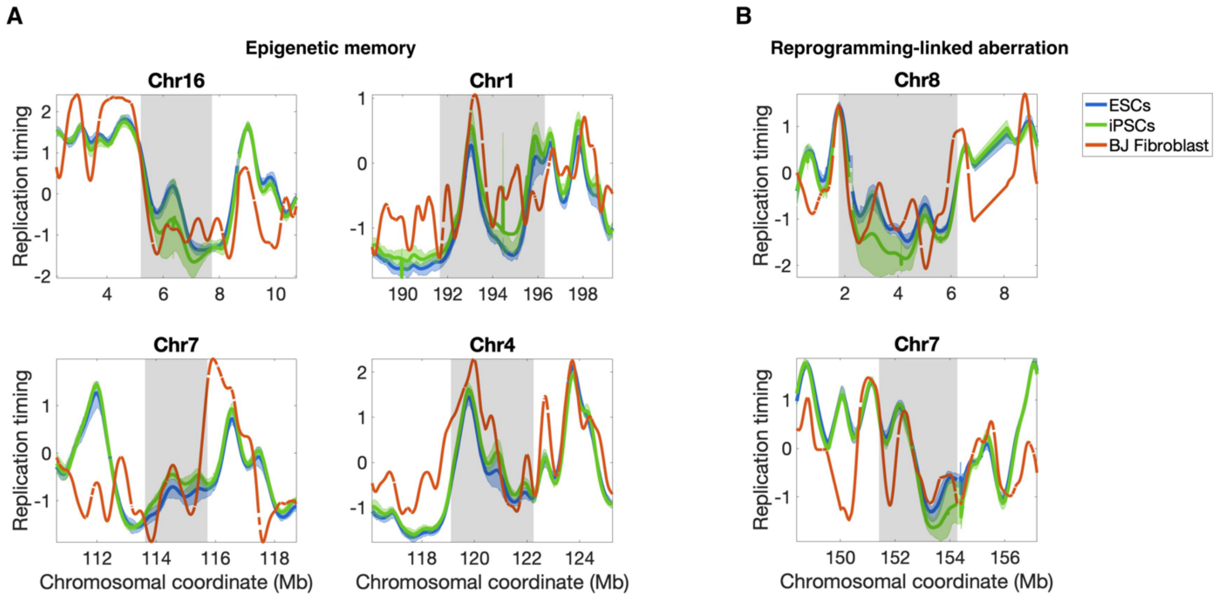


**Supplemental Figure 9**. **Comparing replication timing to fibroblasts suggests both reprogramming-linked and epigenetic memory effects. A**. Fibroblast replication timing derived from single-cells (orange) at four regions inferred to represent epigenetic memory. These regions were defined as those testable regions (**Methods**) where fibroblast replication timing was within +/- 0.2 SD of the iPSC mean (green) at the region of greatest ESC/iPSC variation, and was similarly not within 0.2 SD of the ESC mean (blue). **B**. Fibroblast replication timing at two regions with suspected reprogramming-linked aberration. Regions were defined as in **A**, but with fibroblast replication timing within +/- 0.2 SD of the ESC mean (blue) at the region of greatest ESC/iPSC variation.
